# Tmem117, an oligodendrocyte-enriched regulator of NCX activity, links myelin homeostasis to counterregulation and metabolic health

**DOI:** 10.1101/2025.09.27.678761

**Authors:** Melvin Alappat, Marta Anna Mazurkiewicz, Alexandre Picard, Francesco Prisco, Anja Kipar, Musadiq A Bhat, Dietmar Benke, Hanns Ulrich Zeilhofer, Bernard Thorens, Sevasti Gaspari

**Author notes:** These authors contributed equally (Melvin Alappat, Marta Anna Mazurkiewicz).

## Abstract

The counterregulatory response (CRR) to hypoglycemia is a fundamental, evolutionarily conserved homeostatic mechanism orchestrated by the central nervous system (CNS) to ensure survival during glucose scarcity. In individuals with diabetes, this response is frequently impaired, contributing to life-threatening episodes of hypoglycemia. *Tmem117* was previously identified in a genetic screen as a promising hypothalamic regulator of CRR. Our previous work highlighted its contribution to CRR through regulation of vasopressin secretion.

Here, we reveal that Tmem117 is also enriched in cells of the oligodendrocytic lineage and we characterize the contribution of oligodendrocytic Tmem117 in CRR. We show that depletion of Tmem117 from either all oligodendrocyte lineage cells or only mature oligodendrocytes leads to myelin deficits and male-specific defects in CRR. Furthermore, we reveal that transient, adult-onset depletion of Tmem117 in mature oligodendrocytes is sufficient to induce long-lasting metabolic imbalances in male mice, suggesting that defects in oligodendrocytes and myelin can affect peripheral glucose homeostasis. Mechanistically, we provide for the first-time insights on the function of Tmem117 showing that it regulates intracellular calcium dynamics through its interaction with the sodium-calcium exchanger NCX1.

Together, these results redefine our understanding of the cellular contributors to the CRR, highlight the importance of oligodendrocytes in systemic glucose regulation, and position Tmem117 as a promising molecular target for cell-specific manipulation of NCX activity.

## Intro

Myelin plays a critical role not only in enabling saltatory action potential conduction but also in providing essential trophic support to axons^1^. The integrity of myelin sheaths is maintained by oligodendrocyte lineage cells, which have emerged as key regulators of central nervous system (CNS) homeostasis. Consequently, these cells have become a focal point in research on demyelinating and neurodegenerative diseases^2^. Recent advances have underscored the unique biological features of mature myelinating oligodendrocytes, including their high metabolic demands and particular vulnerability to oxidative stress^3^.

In parallel, accumulating evidence reveals a surprising degree of plasticity within the oligodendroglial lineage. Oligodendrocyte precursor cells (OPCs) and mature oligodendrocytes can dynamically respond to peripheral metabolic and inflammatory cues, suggesting a bidirectional relationship between CNS glial health and systemic physiology^4,5^. However, this relationship has largely been studied in one direction: how peripheral disruptions, such as inflammation or metabolic imbalance, impact oligodendrocyte function and myelin homeostasis. The converse—whether and how oligodendrocyte homeostasis influences whole-body metabolism—remains an open and underexplored question.

In this study, we investigate the role of Tmem117, a poorly characterized transmembrane protein with a strikingly restricted pattern of expression, notably enriched in oligodendrocytes and a few other specialized cell types. Although its function remains unknown, prior work, including our own, has implicated Tmem117 in cellular stress responses^6,7^. Specifically, we previously showed that Tmem117 is required for the survival of vasopressin (AVP)-producing neuroendocrine cells, where its loss leads to ER stress, elevated intracellular calcium levels, and ultimately cell death^7^.

Here, we extend this line of investigation to the oligodendrocyte lineage. Using conditional genetic models, we demonstrate that Tmem117 is necessary for maintaining oligodendrocyte homeostasis and myelin integrity. Loss of Tmem117 in oligodendrocytes leads to sex-specific impairments in the counterregulatory response (CRR) to hypoglycemia, as well as long-term metabolic imbalances.

Notably, we provide *in vitro* evidence that this phenotype arises from disruptions in calcium homeostasis, mediated through a functional interaction between Tmem117 and Slc8a1 (NCX1), a sodium-calcium exchanger. We show that Tmem117 promotes NCX1 stability, thereby facilitating calcium extrusion and preventing cytotoxic calcium overload.

Together, our findings identify Tmem117 as a key regulator of calcium homeostasis, provide mechanistic insight into its function via NCX1 stabilization, and reveal a novel link between CNS myelin health and systemic metabolic regulation.

## Results

### Tmem117 is enriched in oligodendrocytes

*Tmem117* was previously identified in a genetic screening as a candidate gene for the hypothalamic regulation of the CRR to hypoglycemia^8^. Our previous work highlighted its important role on CRR regulation through its effect on AVP neuroendocrine cells^7^. Except from its expression in AVP neuroendocrine cells, publicly available single-cell RNAseq data from the mouse hypothalamus^9^ suggest enrichment of the *Tmem117* transcript in oligodendrocyte lineage cells (Fig 1A, B). The same enrichment is reported also in datasets extending beyond the hypothalamic area (Suppl Fig 1A-C). In the Tabula Muris dataset ^10^ *Tmem117* enriched cells in brain tissue fall within the cluster of oligodendrocytes (Suppl Fig 1A). In addition, data from mousebrain.org^11^ suggest enrichment in mature myelinating oligodendrocytes among others (Suppl Fig 1B,C; highlighted by the red arrow). To test whether the observed enrichment of the transcript is also leading to higher abundance of the Tmem117 protein in oligodendrocytes, we utilized a transgenic mouse line for fluorescent labeling of mature oligodendrocytes and myelin (*CNP-mEGFP*; JAX:026105) alongside immunostaining against Tmem117 using a KO validated antibody^7^ (Fig 1C). Except from the strong immunofluorescence observed in the somata (Fig 1D8,9; magenta arrows) and processes (Fig 1D8-10; turquoise arrows) of AVP neuroendocrine cells, we also observed a fainter but consistent staining in mature oligodendrocytes in the optic nerve (Fig 1D8,9; white arrows). This staining was consistently observed across several brain areas (Fig 1D) with majority of Tmem117 positive cells located in white matter tracts in the form of tightly packed “syncytia” (Fig 1D2,4,6), but some also identified in sparsely myelinated grey matter areas (Fig 1D5,7,10,11). These findings establish that Tmem117 is enriched in mature myelinating oligodendrocytes, suggesting a potential, previously unrecognized, role for this protein in oligodendrocyte biology and myelin homeostasis.

**Figure 1.**
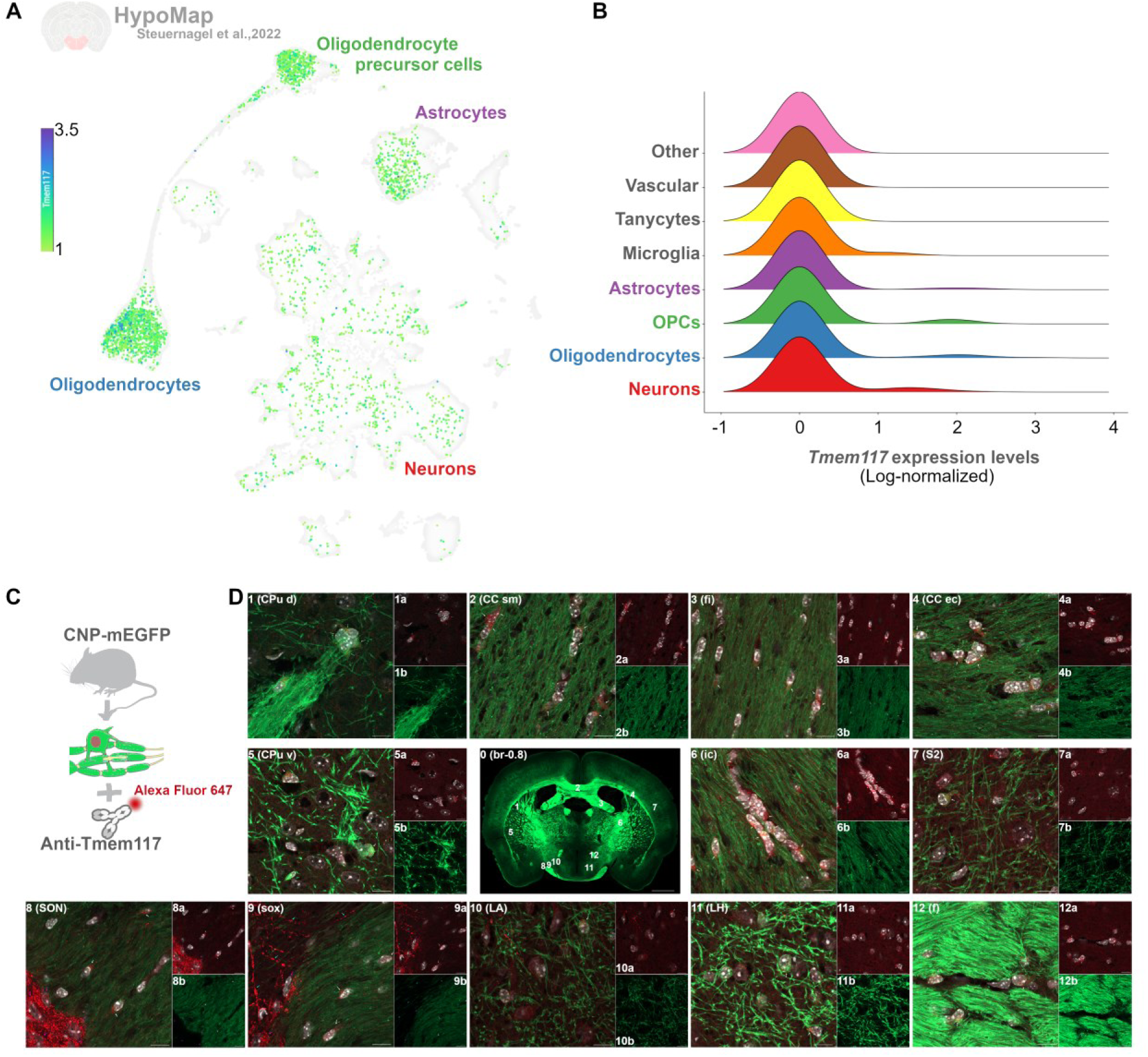
Tmem117 is enriched in oligodendrocytes. **(A)** UMAP depicting the *Tmem117*-enriched cells across all cell-type clusters of the mouse hypothalamus (data extracted from HypoMap). **(B)** Ridge plot depicting the *Tmem117* transcript detection levels across all cell-types in the mouse hypothalamus (data extracted from HypoMap). **(C)** Schematic representation of the approach utilized to verify the detection of Tmem117 in mature oligoderdrocytes. **(D)** Representative images from the mouse brain depicting membrane structures deriving from mature myelinating oligodendrocytes in green (mEGFP) and Tmem117 protein in red (Alexa Fluor 647). Nuclei are labeled with DAPI (white). Magenta and turquoise arrows highlight the somata and processes of AVP neuroendocrine cells, respectively. White arrows highlight Tmem117 positive mature myelinating oligodendrocytes. Scale bars: 1mm for panel 0 and 10μm for panels 1-12, CPu d: striatum dorsal, CC sm: corpus callosum soma, fi: fimbria of hippocampus, CC ec: corpus callosum external capsule, CPu v: striatum ventral, br: bregma, ic: internal capsule, S2: somatosensory cortex, SON: supraoptic nucleus, sox: supraoptic decussation, LA: lateroanterior hypothalamic nucleus, LH: lateral hypothalamic area, f: fornix.

### Depletion of Tmem117 from the oligodendrocyte lineage leads to hypomyelination and male-specific CRR deficits

To study the contribution of oligodendrocytic Tmem117 in CRR we decided to knock-out *Tmem117* in all oligodendrocyte lineage cells by crossing our previously generated *Tmem117 floxed* mouse line^7^ with the Olig2-cre mouse line (JAX:025567) (Fig 2A). Both heterozygote *Tmem117^fl/+^; Olig2-cre^tg/+^* (*Olig2^TM117KO/+^*) and homozygote *Tmem117^fl/fl^; Olig2-cre^tg/+^* (*Olig2^TM117KO/KO^*) male mice were noticeably smaller at weaning and until week 8 of age as reflected by their weight gain curves (Fig 2B, Suppl video 1). In contrast, their female littermates were indistinguishable in size and weight gain when compared to control mice not expressing cre recombinase, *Tmem117^fl/fl^; Olig2-cre^+/+^*(*Olig2^TM117FL/FL^*) (Suppl Fig 2B, Suppl video 1). Furthermore, collective reexamination of the weaning rates over a duration of 26 months (445 pups weaned) revealed decreased representation of *Olig2^TM117KO^*male mice (expected: 50%, observed: 38%) with the rates of female *Olig2^TM117KO^*mice remaining within the expected Mendelian ratio (expected: 50%, observed: 45%; Suppl Fig 2A). When adult (8-week-old) *Olig2^TM117KO^*male mice were subjected to an insulin-induced hypoglycemia (IIH) test (Fig 2C) their CRR appeared diminished with markedly decreased secretion of glucagon (Fig 2D) for hypoglycemic levels comparable to their control *Olig2^TM117FL^* littermates (Fig 2E). In contrast, both hypoglycemia and glucagon secretion were comparable between *Olig2^TM117KO^*and *Olig2^TM117FL^* female mice (Suppl Fig 2C, D). Vagal nerve recordings revealed decreased parasympathetic activity in *Olig2^TM117KO^* male mice during IIH (Fig 2F). Furthermore, transmission electron microscopy analysis of the *Corpus callosum* of *Olig2^TM117KO^* male mice revealed the existence of thinner myelin sheaths (Fig 2G, H) and cytoplasmic inclusions suggestive of myelin instability (Fig 2H). These data demonstrate that *Tmem117* deletion in oligodendrocyte lineage cells impairs myelination and disrupts CRR only in male mice, underscoring a sex-dependent vulnerability to oligodendrocyte dysfunction.

**Figure 2.**
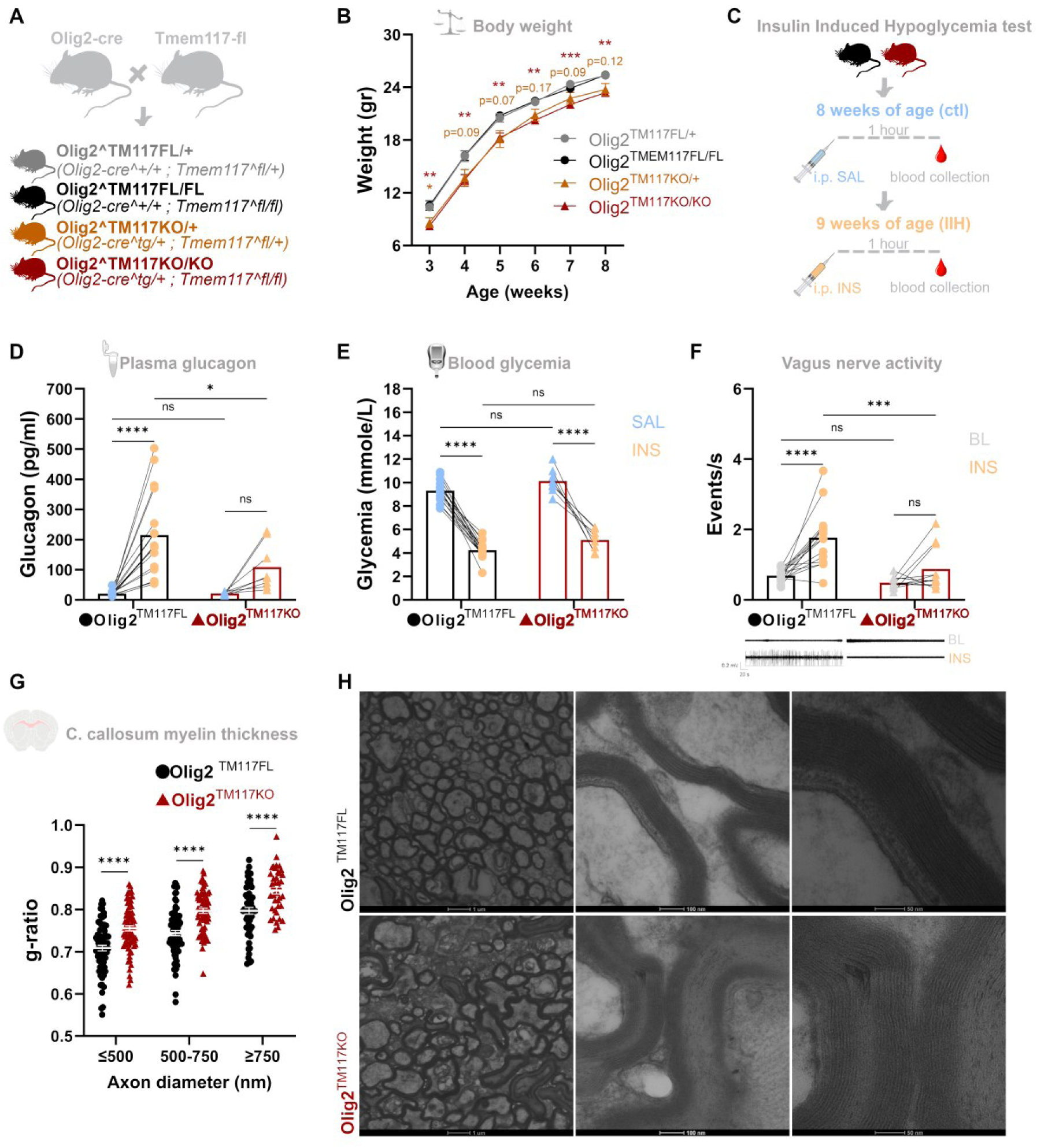
Depletion of Tmem117 from all oligodendrocyte lineage cells in male mice impedes CRR and triggers myelin defects. **(A)** Schematic representation of the transgenic mouse line generation. **(B)** Weight gain curves of male mice missing one (orange; *Olig2^TMEM117KO/+^*) or both (red; *Olig2^TMEM117KO/KO^*) copies of the *Tmem117* gene in oligodendrocyte lineage cells compared to littermate controls (black, gray) with normal expression of *Tmem117* [n=16-29 per group; 2-way ANOVA RM; p values correspond to Dunnett’s multiple comparisons test results for each group when compared to control (*Olig2^TMEM117FL/FL^*) mice]. **(C**) Schematic representation of the insulin induced hypoglycemia paradigm. **(D-E)** Plasma glucagon concentration (D) and blood glucose (E) from *Olig2^TMEM117KO^* male mice (red) and their *Olig2^TMEM117FL^* control littermates (black) one hour after i.p. injection of insulin (yellow) or saline (blue) [n=8-17 per group; 2-way ANOVA RM; p values correspond to Bonferroni’s multiple comparisons test results]. **(F)** Recordings of vagus nerve parasympathetic activity from *Olig2^TMEM117KO^*male mice (red) and their *Olig2^TMEM117FL^* control littermates (black) without any treatment (baseline; gray) or after i.p. injection of insulin (yellow) [n=11-15 per group; 2-way ANOVA RM; p values correspond to Bonferroni’s multiple comparisons test results]. **(G-H)** Electron microscopy analysis of myelin in the *Corpus callosum* of *Olig2^TMEM117KO^* male mice (red) and *Olig2^TMEM117FL^* control littermates (black). G-ratio quantification of myelin thickness (G) and representative images (H) [n=177-209 axons per group corresponding to 2 mice per group; Multiple unpaired t-tests]. Lines/bars correspond to mean values and error bars represent ±SEM. *p<0.05, **p<0.01, ***p<0.001, ****p<0.0001. SAL: saline, INS: insulin, BL: baseline.

### Transient depletion of Tmem117 from mature oligodendrocytes in adult mice leads to demyelination, male specific defects in CRR and long-term metabolic imbalances

Since the male-specific phenotype of the *Olig2^TM117KO^*mouse line could be of developmental origin, we next decided to knock-out *Tmem117* only in mature oligodendrocytes after the critical waves of CNS myelination. To this end, we crossed our *Tmem117 floxed* mouse line^7^ with the Plp1-creER^T^ mouse line (JAX:005975) (Fig 3A). Initially, a group of male *Tmem117^fl/fl^; Plp1-creER^T(tg/+)^*(*Plp1^TM117KO^*) mice and their *Tmem117^fl/fl^; Plp1-creER^T(+/+)^* (*Plp1^TM117FL^*) littermate controls were injected intraperitoneally (i.p.) with tamoxifen at P30-P35 to induce recombination of the *Tmem117* locus in mature oligodendrocytes and then subjected 4 weeks later to the IIH test (Fig 3B). As shown in Fig 3C, male *Plp1^TM117KO^* mice present a CRR phenotype similar to that observed previously in *Olig2^TM117KO^* mice, with diminished glucagon secretion for comparable hypoglycemic states (Fig 3D). Furthermore, transmission electron microscopy analysis of the *Corpus callosum* of male *Plp1^TM117KO^*mice, 4 weeks after induction of recombination by tamoxifen, revealed demyelination and neurodegeneration with high interindividual variability (Fig 3E, each image corresponds to a different mouse). Of note is the myelin phenotype of *Plp1^TM117KO^* mouse number 7 (Fig 3E, bottom right) that from a first overall view appears similar to control mice, but when examined at higher resolution shows subtle signs of myelin decompaction (Fig 3F).

**Figure 3.**
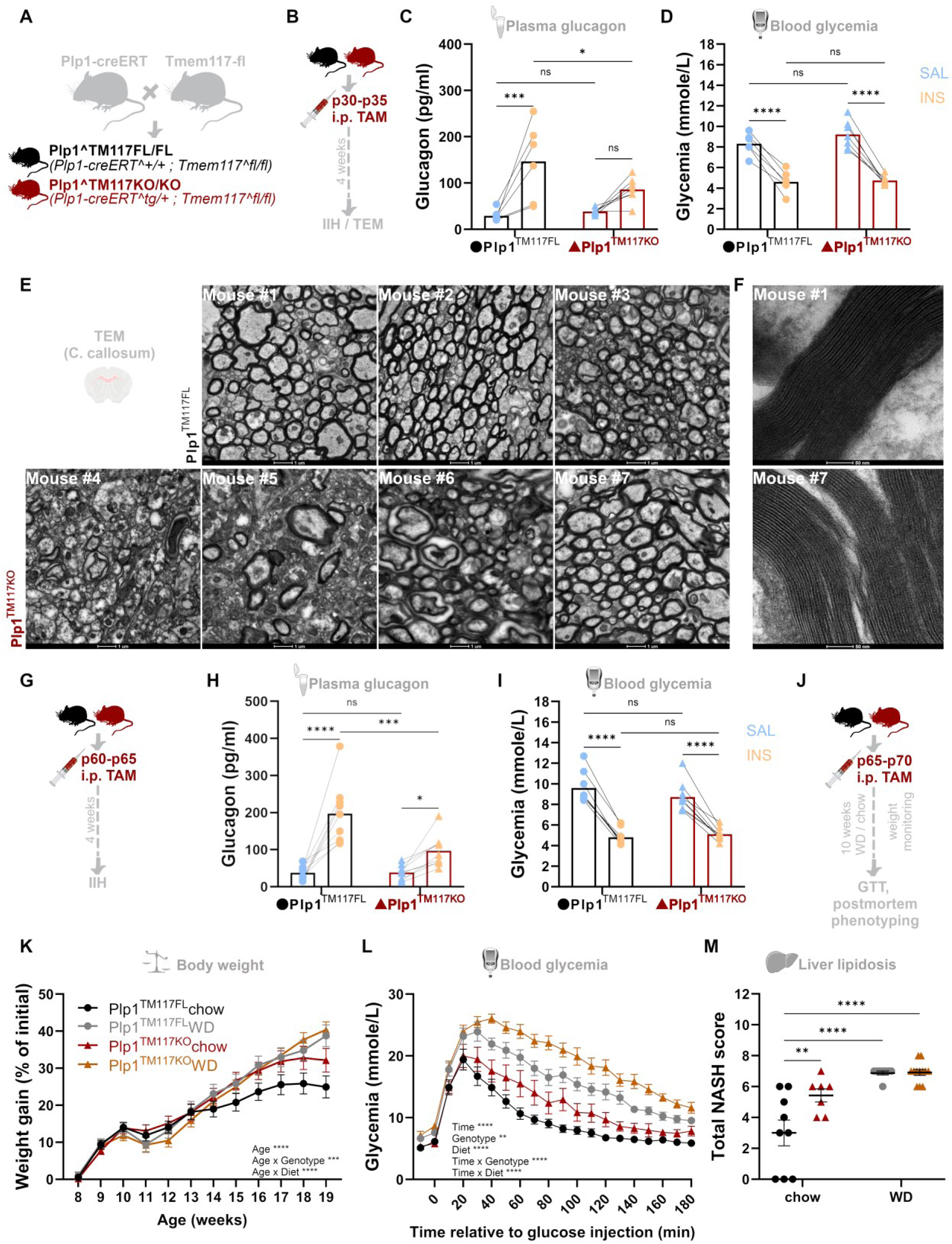
Transient depletion of Tmem117 from mature oligodendrocytes in adult male mice leads to demyelination, defects in CRR and long-term metabolic imbalances. **(A)** Schematic representation of the transgenic mouse line generation. **(B**) Schematic representation of the experimental timeline for panels C-F. **(C-D)** Plasma glucagon concentration (C) and blood glucose (D) from *Plp1^TMEM117KO^*male mice (red) and their *Plp1^TMEM117FL^* control littermates (black) one hour after i.p. injection of insulin (yellow) or saline (blue) [TAM injections performed at P30-P35; n=6-7 per group; 2-way ANOVA RM; p values correspond to Bonferroni’s multiple comparisons test results]. **(E-F)** Representative images of electron microscopy analysis of myelin in the *Corpus callosum* of *Plp1^TMEM117KO^*male mice (botom) and *Plp1^TMEM117FL^* control littermates (top) one month after TAM injection [n=3-4 per group; each image in panel E corresponds to an individual mouse]. **(G)** Schematic representation of the experimental timeline for panels H, I. **(H-I)** Plasma glucagon concentration (H) and blood glucose (I) from *Plp1^TMEM117KO^* male mice (red) and their *Plp1^TMEM117FL^* control littermates (black) one hour after i.p. injection of insulin (yellow) or saline (blue) [TAM injections performed at P60-P65; n=9 per group; 2-way ANOVA RM; p values correspond to Bonferroni’s multiple comparisons test results]. **(J)** Schematic representation of the experimental timeline for panels K-M. **(K)** Weight gain curves of male *Plp1^TMEM117KO^* mice (red, orange) and their *Plp1^TMEM117FL^* control littermates (black, gray) on a regular chow diet (red and black, respectively) or on a WD high in fat and sugar (orange and gray, respectively) [n=10-17 per group; 3-way ANOVA]. **(L)** Glucose tolerance test responses of male *Plp1^TMEM117KO^* mice (red, orange) and their *Plp1^TMEM117FL^* control littermates (black, gray) on a regular chow diet (red and black, respectively) or on a WD high in fat and sugar (orange and gray, respectively) [n=7-13 per group; 3-way ANOVA; week 19 of age]. **(M)** Non-alcoholic steatohepatitis (NASH) scores (mouse-adapted modified Kleiner score) to illustrate the extent of hepatic lipidosis assessed histologically in the liver of male *Plp1^TMEM117KO^* mice (red, orange) and their *Plp1^TMEM117FL^*control littermates (black, gray) on a regular chow diet (red and black, respectively) or on a WD high in fat and sugar (orange and gray, respectively) [n=7-12 per group; 2-way ANOVA; p values correspond to Tukey’s multiple comparisons test results]. Lines/bars correspond to mean values and error bars represent ±SEM. *p<0.05, **p<0.01, ***p<0.001, ****p<0.0001. SAL: saline, INS: insulin, TAM: tamoxifen, WD: western diet.

Then in another group of male mice we induced the recombination in adulthood (P60-65; Fig 3G) and the effect on glucagon secretion remained the same (Fig 3H, I). In contrast, female *Plp1^TM117KO^* mice following exactly the same protocol (P60-65; Suppl Fig3A) responded similar to their *Plp1^TM117FL^* littermate controls (Suppl Fig 3B, C). Of note is that for comparable levels of hypoglycemia (Suppl Fig 3E) female *Plp1^TM117KO^* mice in proestrus secreted higher amounts of glucagon (Suppl Fig 3D), hinting at hormonal modulation of CRR sensitivity.

Given the pronounced effect on CRR we wondered whether the transient depletion of Tmem117 from mature oligodendrocytes could have an effect on the long-term metabolic health of the animals. We generated cohorts of male and female mice with depletion in adulthood (P65-P70; Fig 3J) and monitored their weight gain overtime both under control conditions (chow diet) and upon a metabolic trigger (Western diet (WD) enriched in fat and sugar). As shown in Fig 3K, male *Plp1^TM117KO^*mice on regular chow diet gain more weight compared to their *Plp1^TM117FL^*controls. The same effect but with higher variability is also observed for female *Plp1^TM117KO^* mice (Suppl Fig 3G). A glucose tolerance test performed at the end of the monitoring period (at 19 weeks of age) revealed a male specific effect with male *Plp1^TM117KO^* mice showing higher blood glucose levels in comparison to their *Plp1^TM117FL^* controls, irrespective of the diet (Fig 3L), and female *Plp1^TM117KO^* mice responding identical to their *Plp1^TM117FL^*control littermates (Suppl Fig 3H). A full gross and histological postmortem phenotyping of the different mouse groups did not reveal any pathological changes or suggest functional differences in any organ/tissue except for the liver where differences in the extent of hepatocellular lipidosis were observed. These were further investigated and quantified revealing increased steatosis in the mice that received WD. Furthermore, male *Plp1^TM117KO^* mice on chow diet showed increased lipid accumulation when compared to their *Plp1^TM117FL^* chow controls (Fig. 3M, Suppl Fig4, Suppl table 1). On the contrary, female *Plp1^TM117KO^* chow mice and their *Plp1^TM117FL^* chow controls showed similar levels of lipidosis (Suppl Fig. 3I, Suppl Fig 4, Suppl table 1).

To explore the molecular consequences of Tmem117 loss, we collected the *Corpus callosum* of *Plp1^TM117KO^* male mice and their *Plp1^TM117FL^* littermate controls two weeks after induction of recombination by tamoxifen injection (Fig 4A). Quantitative Real-Time PCR (qRT-PCR) verified the efficient recombination of the *Tmem117* locus (Fig 4B) and showed an increase in spliced XBP1 (sXBP1; Fig 4C) indicative of endoplasmic reticulum (ER)-stress induction. Proteomic analysis identified 483 differentially expressed proteins (410 upregulated, 73 downregulated; Fig. 4D).

**Figure 4.**
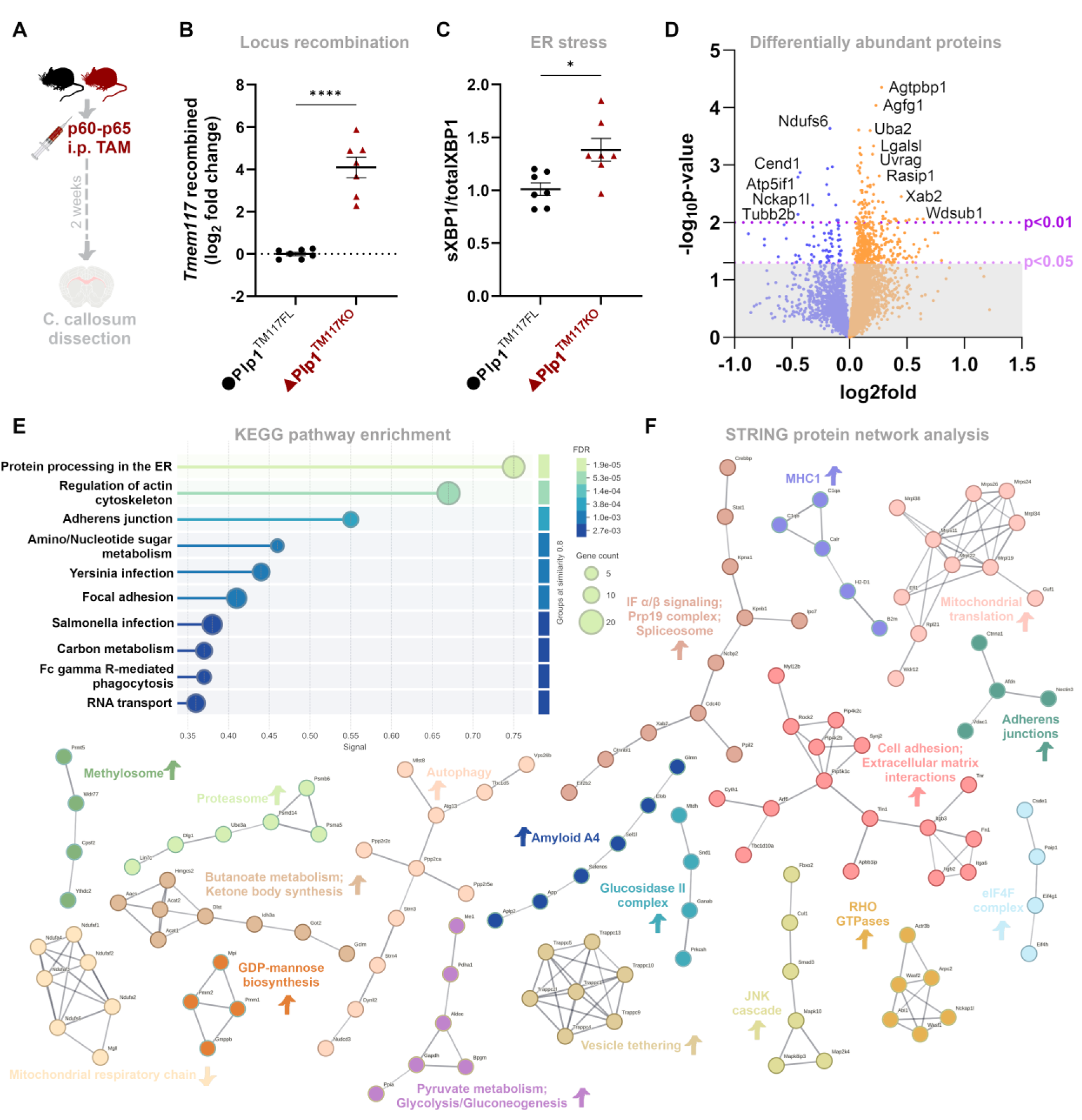
Depletion of Tmem117 from mature oligodendrocytes triggers ER-stress and a wide range of molecular adaptations in the *Corpus callosum*. **(A)** Schematic representation of the experimental timeline. **(B-C)** Real-time PCR results of samples deriving from the *Corpus callosum* of male *Plp1^TMEM117KO^* mice (red) and *Plp1^TMEM117FL^*control littermates (black) verifying the recombination of the *Tmem117* locus (B) and revealing increased levels of spliced *Xbp1* (C) two weeks after TAM injection [n=7 per group; unpaired t-test]. **(D)** Volcano plot depicting the differentially abundant proteins in the *Corpus callosum* of male *Plp1^TMEM117KO^* mice compared to *Plp1^TMEM117FL^*control littermates [n=4 samples per group; 5517 proteins in total; 483 with p<0.05; 90 with p<0.01]. **(E**) STRING KEGG pathway enrichment analysis for all the differentially abundant proteins in the *Plp1^TMEM117KO^ Corpus callosum* that showed a p<0.05. **(F**) STRING functional protein association network analysis with k-means clustering performed for all the differentially abundant proteins in the *Plp1^TMEM117KO^ Corpus callosum* that showed a p<0.05. Lines correspond to mean values and error bars represent ±SEM. *p<0.05, ****p<0.0001. TAM: tamoxifen, IF: interferon, IP: inositol phosphate.

STRING analysis of the 483 differentially expressed proteins revealed that the two most affected pathways in regards to the KEGG pathway database were protein processing in the ER and regulation of actin cytoskeleton (Fig 4E). A closer look to the networks formed by the differentially abundant proteins, focusing only on high-confidence interactions, pointed toward hubs related to respiratory electron transport, mitochondrial translation, vesicle tethering, autophagy, butanoate metabolism, cell adhesion/extracellular matrix, spliceosome/interferon signaling and amyloid formation (Fig 4F).

These results demonstrate that Tmem117 is essential for maintaining oligodendrocyte integrity in adulthood and that its loss induces ER stress and disrupts a wide array of cellular pathways, likely contributing to oligodendrocyte instability and demyelination. Furthermore, they showcase that disruption of oligodendrocyte homeostasis precipitates long-term, sex-specific metabolic dysfunction.

### Tmem117 decreases intracellular calcium by modulating NCX1 activity

The broad cellular disruptions observed, alongside the restricted expression pattern of *Tmem117*, prompted us to ask whether Tmem117 supports homeostasis via a conserved molecular mechanism. Therefore, we decided to look into the specific features of *Tmem117*-enriched cells. From an evolutionary perspective *Tmem117* appears for the first time in vertebrates, but alignment of distant members based on the PFMA motif reveals expression of a TMEM117 protein domain also in cnidaria. To identify the transcriptomic signatures of *Tmem117*-enriched cells across evolution we performed differential expression analysis of *Tmem117*-enriched cells versus cells showing minimal to no expression of *Tmem117* in 3 datasets: cells of the human brain white matter^12^, mouse cells from 20 different organs^10^ and cells of the sea anemone (*Nematostella vectensis*; *v1g11324*-enriched cells)^13^. The results from *M. musculus* and *N. ventensis* were transformed into human orthologs and were overlapped with the results from *H. sapiens*. As expected, there was a very small overlap between all 3 organisms containing only 17 transcripts (Fig 5A). From those 14 ribosomal proteins were commonly downregulated in the *Tmem117*-enriched cells of all organisms, *eno1* was contra regulated (upregulated in *M.musculus* and downregulated in *N.vectensis* and *H.sapiens*) and only 2 transcripts were commonly upregulated: *Tmem117* and *Slc8a1*. *Slc8a1* encodes for NCX1, a sodium-calcium exchanger vital for calcium homeostasis. We had observed in previous studies that Tmem117 depletion led to a strong increase in intracellular calcium in AVP neurons^7^. Therefore, we decided to investigate *in vitro* if overexpression of *Tmem117* in NCX1 expressing cell lines would alter intracellular calcium (Fig 5B). Cells of the mouse hypothalamic neuronal cell line GT1-7 transfected with a plasmid encoding for the mouse *Tmem117* CDS and loaded with the fluorescent calcium indicator Fluo-4 showed lower levels of intracellular calcium upon treatment with KCl compared to non-transfected cells or cells transfected with a mock plasmid not containing the *Tmem117* sequence (Fig 5C). We then performed similar experiments in the mouse pancreatic beta-cell line MIN6B1 to get insight not only into the intracellular calcium levels but also into their effect on the secretory capacity of the cells. Again, similarly to GT1-7 experiments, the cells that were transfected with the *Tmem117* plasmid showed lower intracellular calcium levels upon KCl treatment (Fig 5D). In line with the lower intracellular calcium levels, the insulin secretion of *Tmem117* overexpressing cells was decreased (Suppl Fig 5A). Interestingly the total insulin content of the *Tmem117* overexpressing cells was also decreased (Suppl Fig 5B), an effect that could possibly be explained by a decreased translational activity suggested by our differential expression analysis (Fig 5A). Western blot analysis of the same samples, apart from verifying the overexpression of Tmem117 (Suppl Fig 5C,D), also revealed a negative correlation between the levels of Tmem117 and the amount of insulin secreted (Suppl Fig 5E). To investigate whether this effect on intracellular calcium is mediated through NCX1 we co-transfected HEK293T cells with a plasmid expressing the human TMEM117 tagged with GFP in the C-terminus and a plasmid expressing the human NCX1. The NCX1 plasmid was also carrying a sequence providing resistance over the antibiotic zeocin. Therefore, the cells that survived in a zeocin containing medium and had GFP fluorescence were expressing both proteins of interest. Forty-eight hours after transfection cells were treated with medium containing the ratiometric calcium indicator Fura red alone or in combination with the NCX1 inhibitor SEA0400 for one hour. Then cells were washed, fixed and analyzed for intracellular calcium levels using confocal microscopy. As shown in Fig 5E, a negative correlation between the levels of TMEM117-GFP expression and the concentration of intracellular calcium was observed. Interestingly, this correlation was not present in the cells that were treated with the NCX1 inhibitor (Fig 5F), proving that the lower intracellular calcium observed in Tmem117 overexpressing cells is due to NCX1 activity. Representative images can be found in Fig 5G, with cells showing higher expression of TMEM117-GFP highlighted with white arrows. To check whether this higher NCX1 activity in TMEM117 overexpressing cells is due to higher levels of NCX1 expression we quantified NCX1 levels with immunofluorescence and western blot. Quantification of the NCX1 immunofluorescent analysis revealed a positive correlation between TMEM117-GFP fluorescent intensity and NCX1 fluorescent intensity (Fig 5H, I), suggesting that presence of TMEM117 in higher levels leads to higher levels of NCX1 protein. This suggestion was further proved by western blot analysis that revealed a profound enrichment of NCX1 protein in TMEM117- GFP transfected cells compared to GFP only transfected control cells (Fig 5J, K). Lastly, to investigate if this stabilization of NCX1 by TMEM117 is through a direct interaction we utilized a proximity ligation assay (PLA). As shown in Fig 5L and M, the PLA assay revealed a strong signal in TMEM117-GFP expressing cells that was not present in GFP only transfected control cells.

**Figure 5.**
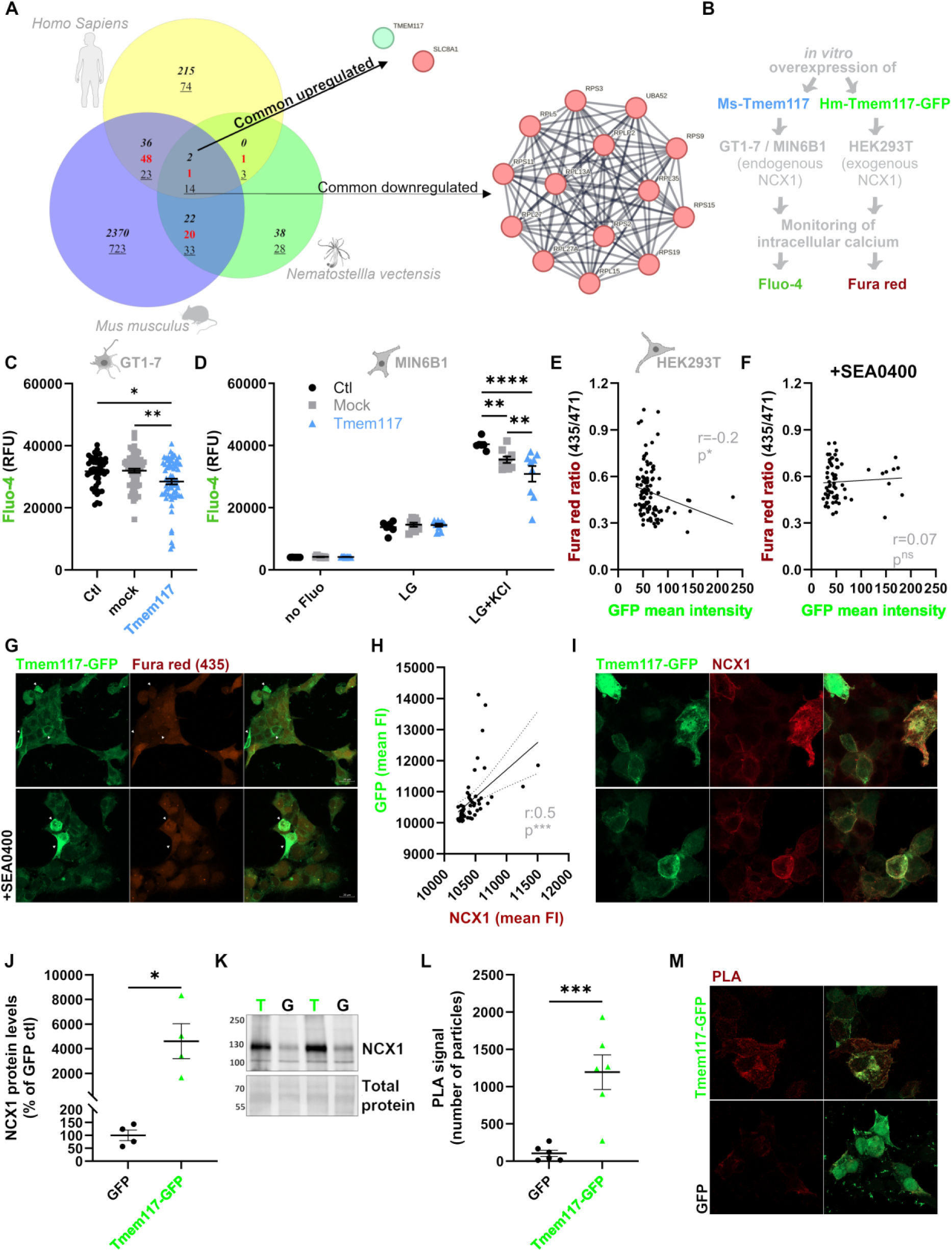
Tmem117 decreases intracellular calcium by modulating NCX1 activity. **(A)** Venn diagram depicting the overlap of the differentially expressed genes in *Tmem117*-enriched cells compared to cells that express low to no *Tmem117* across 3 evolutionary distinct organisms (bold: common upregulated, red: contra-regulated, underlined: common downregulated). **(B)** Schematic representation of the experimental outline. **(C)** Quantification of intracellular calcium in GT1-7 cells expressing *Tmem117* (blue) versus mock transfected (gray) or non-transfected (black) cells [n=48-66 wells per group; 1-way ANOVA; p values correspond to Tukey’s multiple comparisons test results]. **(D)** Quantification of intracellular calcium in MIN6B1 cells expressing *Tmem117* (blue) versus mock transfected (gray) or non-transfected (black) cells [n=6-9 wells per group; 2-way ANOVA; p values correspond to Tukey’s multiple comparisons test results]. **(E-G)** Ratiometric quantification of intracellular calcium in HEK293T cells co-expressing TMEM117-GFP and NCX1. Correlation of the GFP intensity (used as a proxy for TMEM117 expression levels) with intracellular calcium without (E) or with (F) the NCX1 inhibitor SEA0400. Representative images (G) with cells showing higher expression of TMEM117-GFP highlighted by the white arrows [n=68-108 cells per group; Pearson’s r correlation analysis]. **(H-I)** Immunofluorescent analysis of HEK293T cells co-expressing TMEM117-GFP and NCX1. Correlation of the GFP intensity (used as a proxy for TMEM117 expression levels) with the intensity of NCX1 staining (H). Representative images (I) [n=52 cells; Pearson’s r correlation analysis]. **(J-K)** Western blot quantification (J) and representative blot (K) of HEK293T cells co-expressing GFP and NCX1 or TMEM117-GFP and NCX1 [n=4 per group; unpaired t-test]. **(L-M)** Proximity ligation assay quantification (L) and representative image (M) of HEK293T cells co-expressing GFP and NCX1 or TMEM117-GFP and NCX1 [n=6 per group; unpaired t-test].

These results identify TMEM117 as a molecular stabilizer of NCX1, suggesting a unifying mechanism by which TMEM117 maintains calcium homeostasis and supports cell viability in specialized cell types such as oligodendrocytes and neuroendocrine cells.

## Discussion

*Tmem117* was initially identified in a genetic screen that aimed at uncovering hypothalamic regulators of the CRR^8^. In our previous work, we demonstrated its functional relevance in AVP-producing neuroendocrine cells^7^. Here, by mining publicly available RNAseq datasets, we expand on these findings, identifying an enrichment of *Tmem117* in cells of the oligodendrocyte lineage and establishing its essential role in maintaining myelin homeostasis.

Closer inspection of single-cell RNAseq data revealed that although *Tmem117* is particularly enriched in oligodendrocytes, it is also expressed in distinct subpopulations of neurons and hypothalamic astrocytes (Fig. 1A, C). Beyond the central nervous system, analysis of the Tabula Muris dataset revealed a similarly selective expression profile in peripheral tissues, including subsets of large intestine epithelial cells, cardiomyocytes, and kidney collecting duct cells. This cell-type specificity prompted us to perform a differential expression analysis comparing *Tmem117*-enriched cells to non-enriched populations. Applying an evolutionary filter to this analysis led to the identification of *Slc8a1*, which encodes the sodium-calcium exchanger NCX1, as the only transcript consistently co-enriched across all *Tmem117*-enriched cells. For these cross-species comparisons, we mapped *M. musculus* and *N. vectensis* transcripts to their *H. sapiens* orthologs. Consequently, while *Slc8a1* emerged as the main candidate, its paralogs *Slc8a2* (NCX2) and *Slc8a3* (NCX3) are also orthologs of the *N. vectensis* transcript *v1g239709*, which was enriched in cells expressing *v1g11324* (the gene encoding the TMEM117 protein domain). Although we focus here on *in vitro* validation of the TMEM117–NCX1 interaction, our data raise the possibility that TMEM117 also modulates NCX2 and NCX3 function. Given the known importance of NCX3 in mature oligodendrocytes^14^, its potential involvement—alongside NCX1—in the observed phenotypes becomes a priority for future studies.

Our *in vitro* findings show that TMEM117 stabilizes NCX1 and modulates intracellular calcium dynamics, pointing to a regulatory mechanism with broad physiological relevance. Calcium homeostasis is fundamental to cellular function, and NCX activity plays a pivotal role in numerous pathological contexts^15^. Notably, a recent study reported that *Tmem117* knockdown is protective against cardiac hypertrophy^16^—a condition in which NCX1 is upregulated. Our data suggest a mechanistic basis for this observation: TMEM117-dependent stabilization of NCX1. In oligodendrocytes, NCX1 and NCX3 are differentially regulated during lineage progression—NCX1 is predominant in OPCs and downregulated with maturation, whereas NCX3 follows the opposite pattern^14^. The enrichment of *Tmem117* in both OPCs and mature oligodendrocytes (Fig. 1A) supports the hypothesis that it may serve as a stage-specific modulator of calcium signaling via these exchangers.

Apart from molecular interactions, our study highlights a broader physiological principle: oligodendrocyte homeostasis is critical—particularly in males—for CRR and systemic metabolic regulation. We demonstrate that *Tmem117* deletion throughout the oligodendrocyte lineage impairs CRR and causes myelin abnormalities in male mice. Strikingly, inducible deletion of *Tmem117* in mature oligodendrocytes during adulthood recapitulates these effects and leads to long-term alterations in metabolic profiles. This suggests that even transient disruptions in mature oligodendrocyte function—despite the capacity for OPC-mediated remyelination—can result in enduring physiological consequences. CRR has traditionally been considered a strictly neuronal process. Glucose-sensing neurons in key brain regions detect fluctuations in blood glucose and trigger neuroendocrine responses to restore homeostasis^17^. Our results suggest that non-neuronal cells, specifically oligodendrocytes, also play a critical role—likely by influencing axonal conduction and signal fidelity within this circuitry. Furthermore, our findings related to weight gain, glucose tolerance, and hepatic lipid deposition demonstrate for the first time that oligodendrocyte dysfunction alone can impact whole-body metabolism. To date, most studies have adopted a bottom-up perspective, examining how systemic metabolic imbalances affect CNS cell populations^18^. Our data introduce a complementary, top-down model in which oligodendrocyte dysfunction and disrupted myelin integrity act as upstream drivers of systemic metabolic imbalance. Importantly, these effects were sexually dimorphic. We observed them exclusively in male mice, while female mice exhibited either no phenotype or, intriguingly, enhanced CRR during proestrus (Fig. 3G). This points toward a role for sex hormones in modulating central glucose sensing and neuroglial interactions. Previous studies have reported sex differences in oligodendrocyte biology and myelin dynamics^19–21^, and sexual dimorphism in systemic metabolism is well documented^22^. Whether the observed sex-specific effects are due to intrinsic differences in the oligodendrocyte lineage or reflect sexually dimorphic regulation of *Tmem117* itself remains an open question for future investigation.

Beyond *Tmem117*’s sexual dimorphism, several additional questions arise from our findings. Electron microscopy analysis of the Corpus callosum suggests that our *Plp1^TMEM117KO^* mouse line may represent a novel model for inducible demyelination. However, more extensive characterization across the CNS is required to confirm this. Because a full survey of demyelination would be challenging with traditional TEM, we are now crossing our *Plp1^TMEM117KO^* mice with the Tg(Cnp-EGFP)1Qrlu/J* line. This will allow us to use clearing and light-sheet microscopy for whole-CNS analysis of myelination, enabling us to assess both spatial and temporal dynamics of demyelination and remyelination *in vivo*. In parallel, we are also exploring tools to directly assess how Tmem117 influences NCX1 and NCX3 in oligodendrocytes *in vivo*—an effort that could yield valuable insights for targeting demyelinating diseases. Finally, given *Tmem117*’s restricted expression in tissues such as the heart, kidney, and colon, we hope that our mechanistic insights into NCX1 regulation will inspire further investigation into its role in other organ systems and disease contexts.

In conclusion, our study identifies Tmem117 as a novel modulator of NCX1 activity with highly cell-specific expression and establishes oligodendrocyte homeostasis as a key regulator of systemic metabolic health. These findings open the door to further mechanistic dissection and suggest new therapeutic avenues for diseases involving myelin dysfunction and calcium signaling dysregulation.

## Acknowledgements

We are grateful to Christel Genoud and Jean Daraspe from the Electron Microscopy Facility (EMF) of University of Lausanne for transmission electron microscopy sample processing and imaging and to the Protein Analysis Facility (PAF) of University of Lausanne for the proteomic analysis of the *Corpus callosum* samples. We are also grateful to Dr. Simon Quenneville for cloning the Tmem117 CDS in the pRRL-PGK-Tmem117-flag-puroR plasmid. We would also like to thank Dr. Arne Battefeld for kindly providing us with the Tg(Cnp-EGFP*)1Qrlu/J mouse line and Prof. G Van der Goot for kindly providing the TMEM117-GFP plasmid. We also wish to thank the lab technician team of the Histology Laboratory, Institute of Veterinary Pathology, Vetsuisse Faculty, University of Zurich, for excellent technical support. Furthermore, we are grateful to Dr. Nicolas Geux for aligning distant members based on the PFMA motif of the TMEM117 domain and for his valuable scientific insights into unraveling the function of Tmem117. We would also like to thank Dr. Lukas Steuernagel for generating the ridge plot of *Tmem117* expression in the HypoMap dataset. Lastly, we would like to thank Prof. Aiman Saab for his valuable input on the manuscript. This work was supported by a European Research Council Advanced Grant (Integrate, No. 694798) and a Swiss National Science Foundation grant (310030-182496) to BT and a Swiss National Science Foundation Ambizione grant (208875) and a Postdoc grant from University of Zurich (K-41601-02-01) to SG. Illustrations were adapted from NIAID NIH BIOART Source (bioart.niaid.nih.gov/bioart/).

## Author Contributions

**MA** formal analysis; investigation; visualization; methodology; writing – review and editing. **MAM** formal analysis; investigation; visualization; methodology; writing – review and editing. **AP** formal analysis; investigation. **FP** formal analysis; investigation. **AK** formal analysis; investigation; writing – review and editing. **MAB** methodology; resources. **DB** methodology; resources. **HUZ** resources; supervision. **BT** conceptualization; resources; supervision; funding acquisition; writing – review and editing. **SG** conceptualization; formal analysis; investigation; visualization; methodology; writing –original draft, review and editing; project administration; funding acquisition.

## RESOURCE AVAILABILITY

### Lead contact

Further information and requests for resources and reagents should be directed to and will be fulfilled by the Lead Contact, Sevasti Gaspari.

### Materials availability

Free of charge for non-commercial purposes upon request to the lead author.

### Data availability

Data reported in this paper will be shared by the lead contact upon request.

Any additional information required to reanalyze the data reported in this paper is available from the lead contact upon request.

All raw MS data together with raw output tables are available via the Proteomexchange data repository (www.proteomexchange.org) with the accession PXD064620.

## Code availability

The original code used for the DEA of the transcriptomic datasets is available from the lead contact upon request.

## DECLARATION OF GENERATIVE AI AND AI-ASSISTED TECHNOLOGIES IN THE WRITING PROCESS

During the preparation of this work the authors used Microsoft Copilot and ChatGPT exclusively for text polishing as well as language and grammar corrections. After using these tools, the authors reviewed and edited the content as needed and take full responsibility for the content of the publication.

---

**Supplemental Figure 1.**
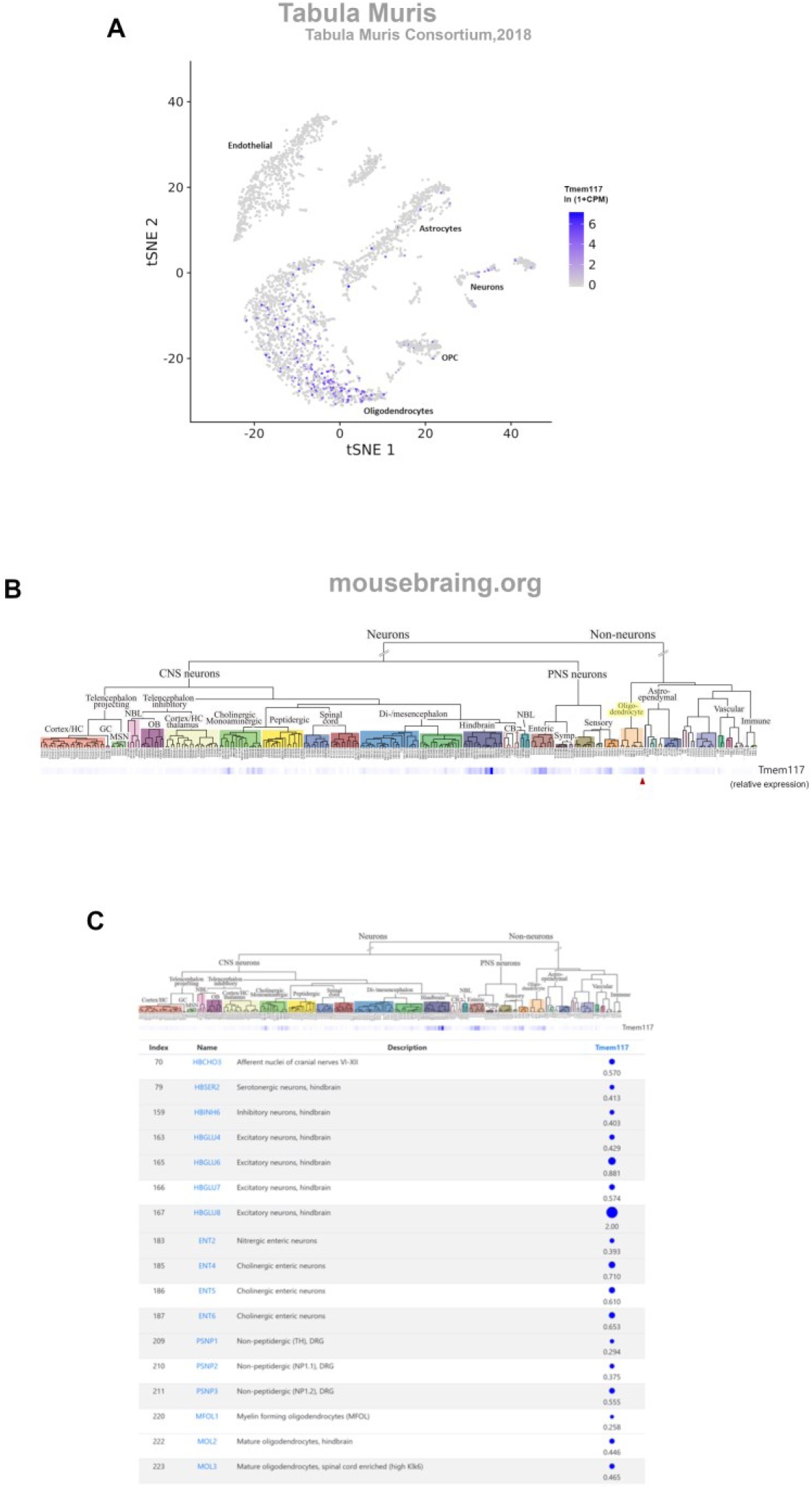
Tmem117 is enriched in oligodendrocytes. **(A)** UMAP depicting the *Tmem117* transcript detection levels across different cell-types in the mouse brain (data extracted from Tabula Muris). **(B)** Dendrogram depicting the *Tmem117* transcript detection levels across different cell-types in the mouse nervous system (data extracted from mousebrain.org). Mature myelinating oligodendrocytes are highlighted by the red arrow. **(C)** Depiction of the *Tmem117*-expressing cell clusters and its relative levels of expression in each one of them (data extracted from mousebrain.org).

**Supplemental Figure 2.**
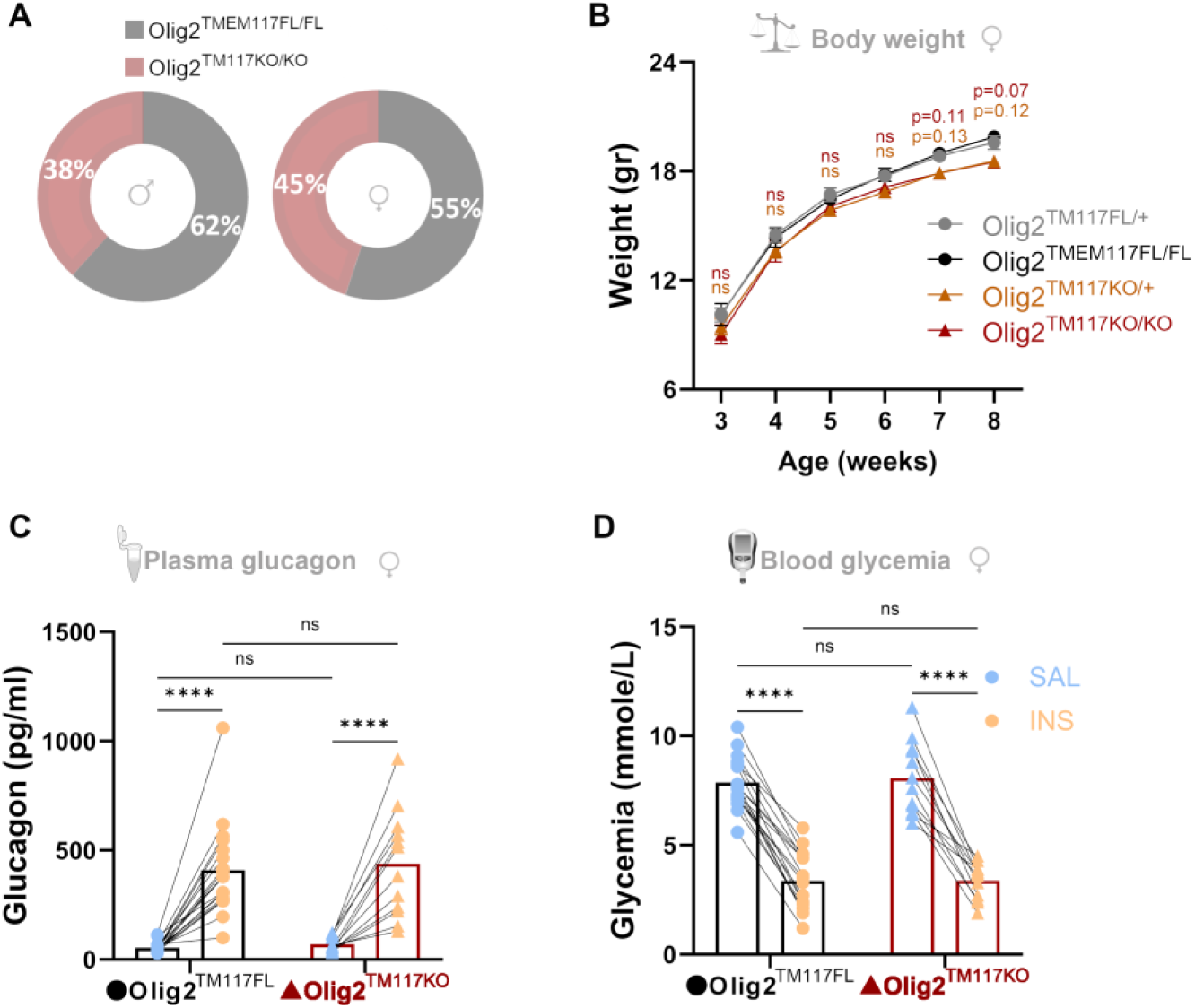
Depletion of Tmem117 from all oligodendrocyte lineage cells in female mice. **(A)** Weaning rates of male (left) and female (right) *Olig2^TMEM117KO/KO^* mice (red) and their *Olig2^TMEM117FL/FL^* control littermates (black). The expected rate based on mendelian genetics is 50% [n=445 pups weaned within a period of 26 months; breeding scheme: *Tmem117^fl/fl^; Olig2cre^tg/+^* x *Tmem117^fl/fl^; Olig2cre**^+/+^***]. **(B)** Weight gain curves of female mice missing one (orange; *Olig2^TMEM117KO/+^*) or both (red; *Olig2^TMEM117KO/KO^*) copies of the *Tmem117* gene in oligodendrocyte lineage cells compared to littermate controls (black, gray) with normal expression of *Tmem117* [n=19-29 per group; 2-way ANOVA RM; p values correspond to Dunnett’s multiple comparisons test results for each group when compared to control (*Olig2^TMEM117FL/FL^*) mice]. **(C-D)** Plasma glucagon concentration (C) and blood glucose (D) from *Olig2^TMEM117KO^* female mice (red) and their *Olig2^TMEM117FL^* control littermates (black) one hour after i.p. injection of insulin (yellow) or saline (blue) [n=12-18 per group; 2-way ANOVA RM; p values correspond to Bonferroni’s multiple comparisons test results]. Lines/bars correspond to mean values and error bars represent ±SEM. *p<0.05, **p<0.01, ***p<0.001, ****p<0.0001. SAL: saline, INS: insulin.

**Supplemental Figure 3.**
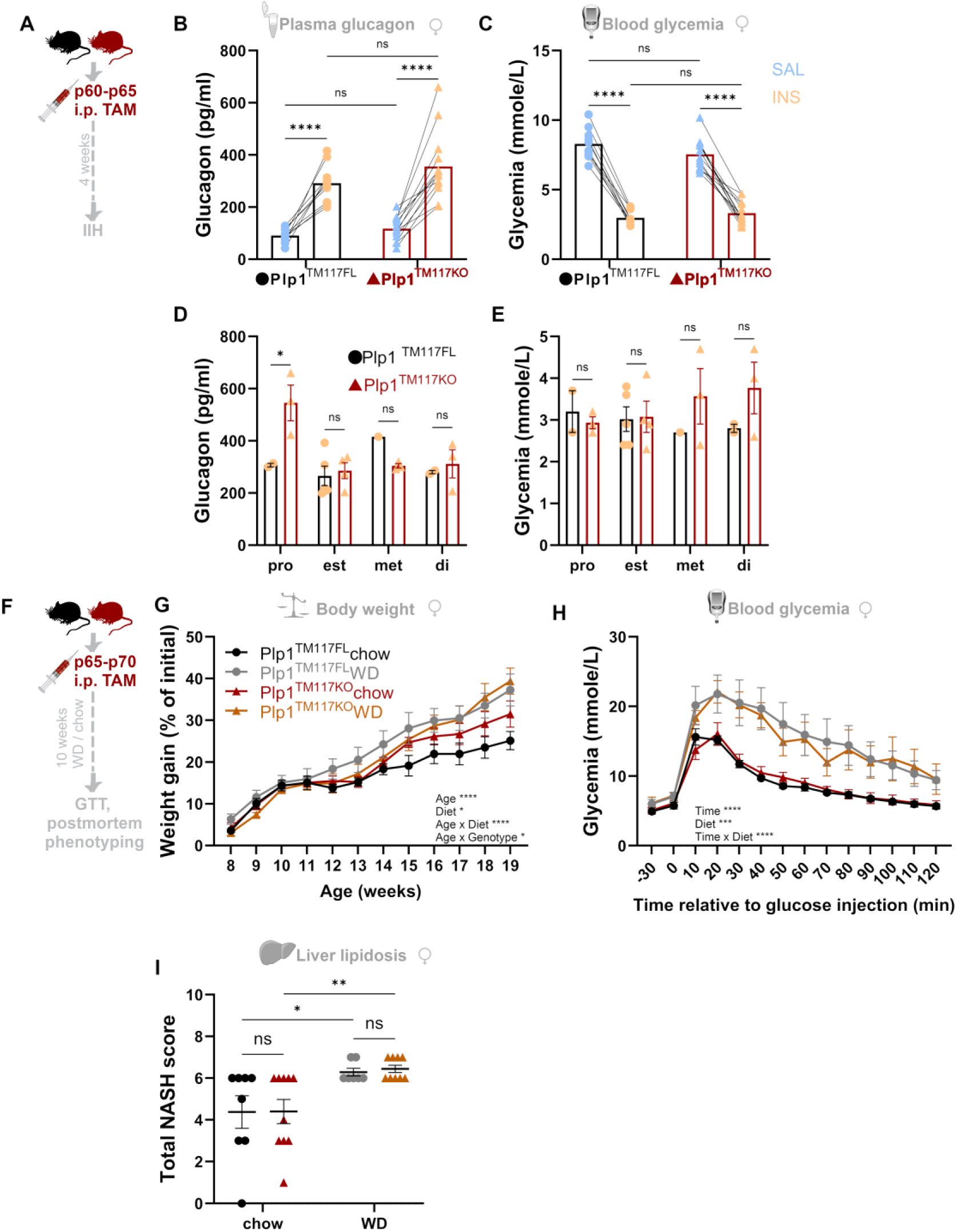
Transient depletion of Tmem117 from mature oligodendrocytes in adult female mice. **(A)** Schematic representation of the experimental timeline for panels B-E. **(B-C**) Plasma glucagon concentration (B) and blood glucose (C) from *Plp1^TMEM117KO^* female mice (red) and their *Plp1^TMEM117FL^* control littermates (black) one hour after i.p. injection of insulin (yellow) or saline (blue) [TAM injections performed at P60-P65; n=10-13 per group; 2-way ANOVA RM; p values correspond to Bonferroni’s multiple comparisons test results]. **(D-E)** Plasma glucagon concentration (D) and blood glucose (E) from *Plp1^TMEM117KO^* female mice (red) and their *Plp1^TMEM117FL^*control littermates (black) one hour after i.p. injection of insulin separated by the phase of the oestrous cycle [the same data as on the INS points of panels B and C; n=10-13 per group; 2-way ANOVA; p values correspond to Bonferroni’s multiple comparisons test results]. **(F)** Schematic representation of the experimental timeline for panels G-I. **(G)** Weight gain curves of female *Plp1^TMEM117KO^* mice (red, orange) and their *Plp1^TMEM117FL^* control littermates (black, gray) on a regular chow diet (red and black, respectively) or on a WD high in fat and sugar (orange and gray, respectively) [n=9-12 per group; 3-way ANOVA]. **(H)** Glucose tolerance test responses of female *Plp1^TMEM117KO^*mice (red, orange) and their *Plp1^TMEM117FL^* control littermates (black, gray) on a regular chow diet (red and black, respectively) or on a WD high in fat and sugar (orange and gray, respectively) [n=5-7 per group; 3-way ANOVA; week 19 of age]. **(I)** Non-alcoholic steatohepatitis (NASH) scores (mouse-adapted modified Kleiner score) to illustrate the extent of hepatic lipidosis assessed histologically in the liver of female *Plp1^TMEM117KO^* mice (red, orange) and their *Plp1^TMEM117FL^* control littermates (black, gray) on a regular chow diet (red and black, respectively) or on a WD high in fat and sugar (orange and gray, respectively) [n=7-10 per group; 2-way ANOVA; p values correspond to Tukey’s multiple comparisons test results]. Lines/bars correspond to mean values and error bars represent ±SEM. *p<0.05, **p<0.01, ***p<0.001, ****p<0.0001. SAL: saline, INS: insulin, pro: proestrus, est: estrus, met: metestrus, di: diestrus, TAM: tamoxifen, WD: western diet.

**Supplemental Figure 4.**
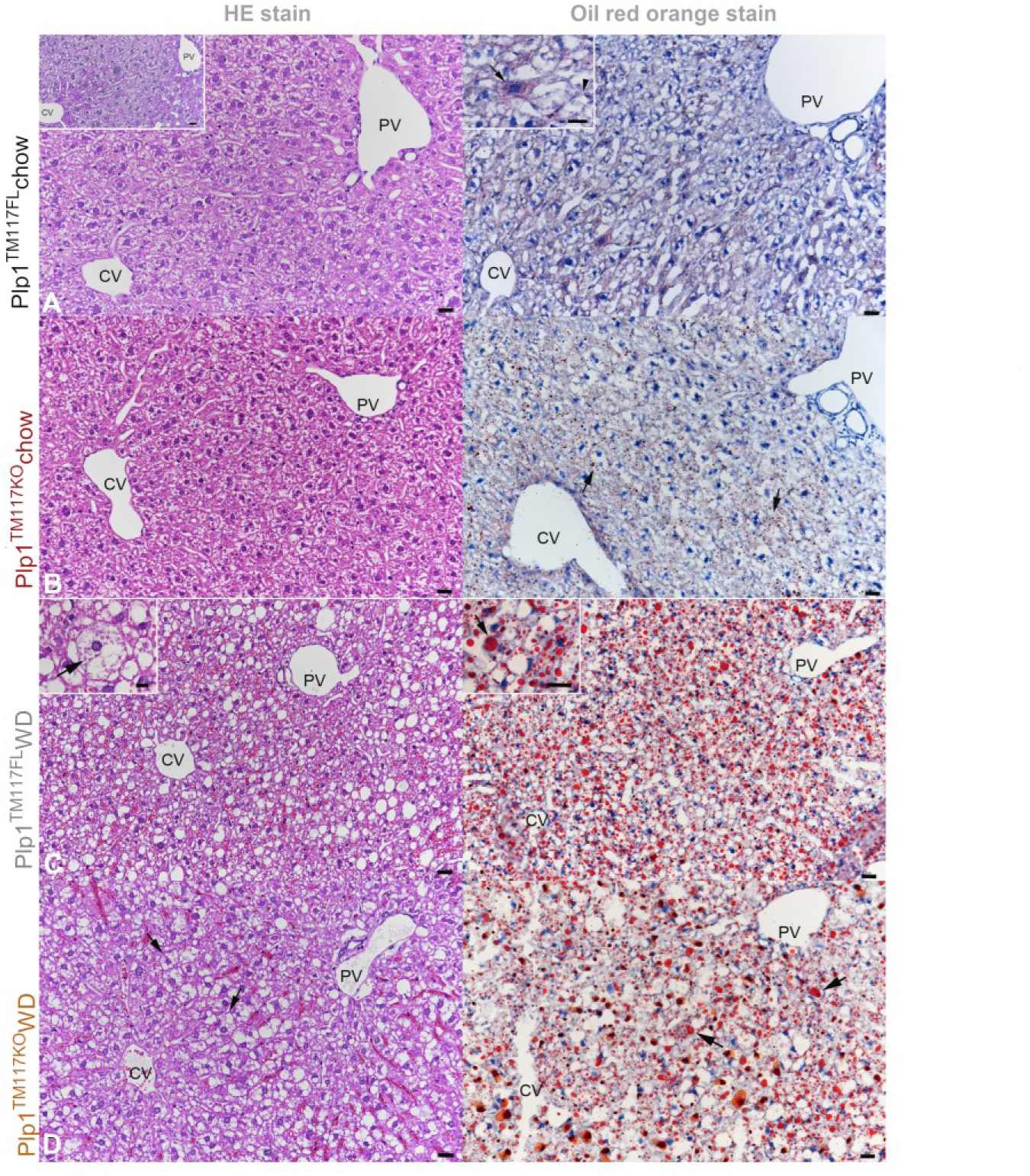
Histological features and extent of hepatocellular steatosis in male *Plp1^TMEM117KO^*mice and *Plp1^TMEM117FL^* controls after chow or WD diet. Left column: HE stains; right column: oil red orange stain to confirm fat deposition; bars = 25 µm. **(A)** *Plp1^TMEM117FL^* control mouse on chow diet, score 3m. Hepatocytes exhibit cloudy vacuolation due to glycogen accumulation (inset left: PAS reaction). The fat stain on the right highlights the presence of microvesicular steatosis, represented by individual (inset: arrowhead) to several (inset: arrow) small fat vacuoles in the cytoplasm of less than a third of hepatocytes. Bar (insets): 25 µm (left), 20 µm (right). **(B)** *Plp1^TMEM117KO^* mouse on chow diet, score 5m. Hepatocytes exhibit cloudy vacuolation due to glycogen accumulation. The fat stain on the right highlights the presence of microvesicular steatosis (arrows) in more than a half of the hepatocytes. **(C)** *Plp1^TMEM117FL^*control mouse on WD, score 7B. A large proportion of hepatocytes exhibit well defined cytoplasmic vacuoles (left image) which represent large fat vacuoles (right) present within the cells together with small fat vacuoles (inset: arrow), representing macrovesicular steatosis. The inset on the left shows a swollen, fat laden hepatocyte (arrow) in early ballooing degeneration. Bars (insets): 10 µm (left), 20 µm (right). **(D)** *Plp1^TMEM117KO^* mouse on WD, score 8BB. A large proportion of hepatocytes exhibit well defined cytoplasmic vacuoles (left image) with several swollen hepatocytes (early ballooing degeneration; arrows). The fat stain (left image) highlights micro- and macrovesicular steatosis of almost all hepatocytes. CV: central vein; PV: portal vein within portal area.

**Supplemental Figure 5.**
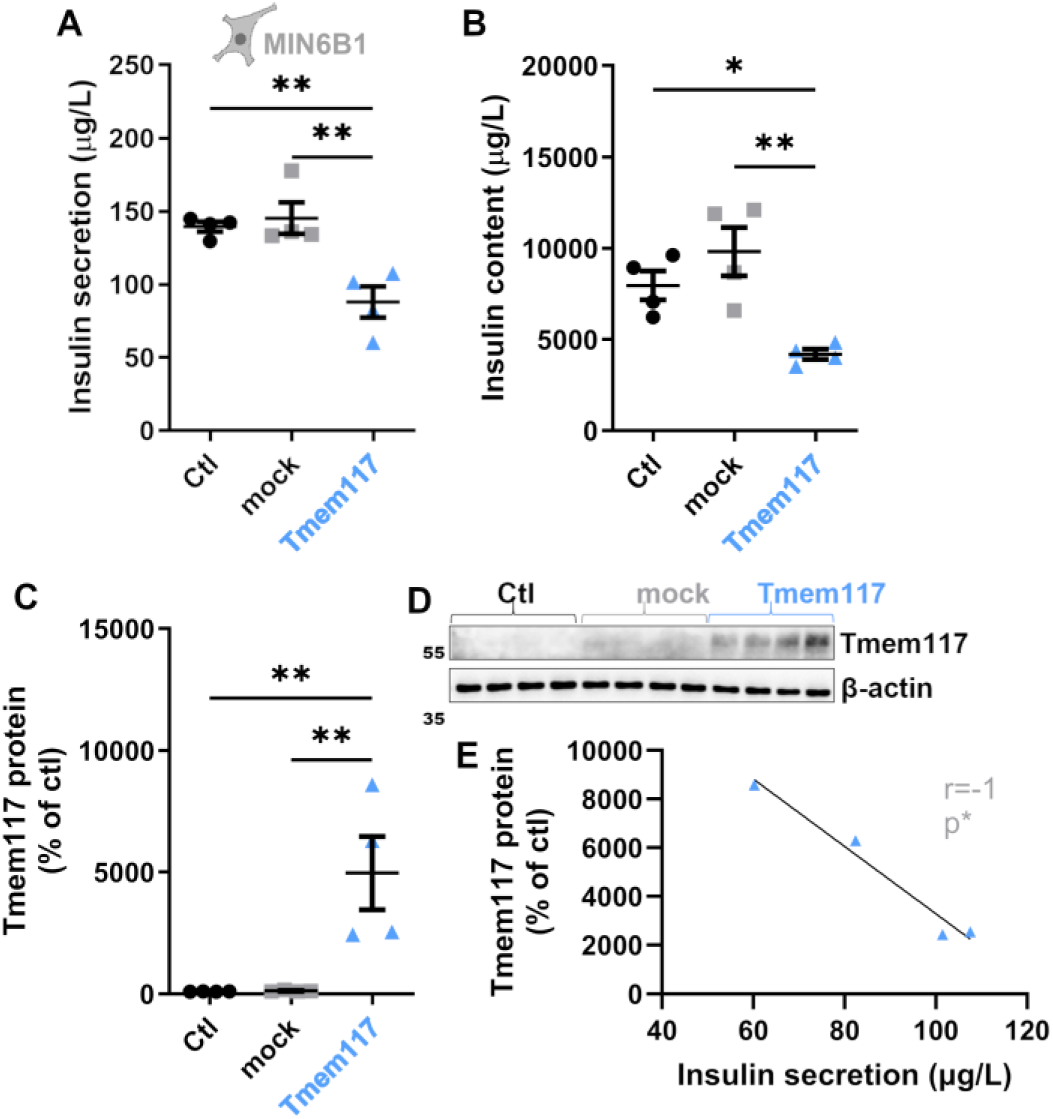
Tmem117 overexpression in MIN6B1 cells decreases insulin content and secretion. **(A-B)** Quantification of insulin secretion (A) and content (B) of MIN6B1 cells expressing *Tmem117* (blue) versus mock transfected (gray) or non-transfected (black) cells [n=4 wells per group; 1-way ANOVA; p values correspond to Tukey’s multiple comparisons test results]. **(C-E)** Western blot quantification (C) and blot (D) of MIN6B1 cells expressing *Tmem117* (blue) versus mock transfected (gray) or non-transfected (black) cells [n=4 wells per group; 1-way ANOVA; p values correspond to Tukey’s multiple comparisons test results]. Correlation analysis (E) between the levels of Tmem117 protein and the amount of secreted insulin [n=4; Pearson’s r correlation analysis].

**Supplemental Table 1.** Hepatic lipidosis scores for each individual animal. Mouse adapted, modified Kleiner scores of individual male and female *Plp1^TM117FL^* and *Plp1^TM117KO^*mice, aged 19 weeks that had been maintained on a regular chow diet (Chow) or a Western Diet enriched in fat and sugar (WD) for 10 weeks. M: exclusively microvesicular steatosis; m+: microvesicular, with occasional medium sized fat vacuoles; m++: microvesicular, with occasional up to macrovesicular; B: scattered/occasional hepatocytes with evidence of ballooning degeneration; BB: many hepatocytes with evidence of ballooning degeneration.

|  | *Plp1*<sup>TM117FL</sup> Chow | *Plp1*<sup>TM117FL</sup> WD | *Plp1*<sup>TM117KO</sup> Chow | *Plp1*<sup>TM117KO</sup> WD |
| --- | --- | --- | --- | --- |
| Male | 3m | 7B | 5m | 7B |
|  | 0 | 7B | 4m | 7B |
|  | 0 | 7B | 4m | 7B |
|  | 4m | 7B | 7m+ | 8BB |
|  | 3m | 7B | 6m+ | 8BB |
|  | 0 | 7B | 6m+ | 7B |
|  | 6m+ | 7 | 6m+ | 7 |
|  | 6m+ | 6 |  | 7B |
|  | 5m |  |  | 6 |
|  |  |  |  | 6 |
|  |  |  |  | 6 |
|  |  |  |  | 7B |
| Female | 6m | 6 | 6m | 6 |
|  | 0 | 6 | 6m | 7B |
|  | 3m | 7B | 1 | 7B |
|  | 3m | 6 | 3m | 6 |
|  | 6m | 7 | 3m | 6 |
|  | 6m+ | 6 | 6m | 6 |
|  | 6m | 6 | 6m | 7 |
|  | 5m |  | 6m+ | 7 |
|  |  |  | 3m | 6 |
|  |  |  | 4m |  |

## Materials & Methods

### EXPERIMENTAL MODELS AND SUBJECT DETAILS

#### Mice

All animal care and experimental procedures were in accordance with the Swiss National Institutional Guidelines of Animal Experimentation (OExA; 455.163) with license-approval (VD3363, VD3674, ZH35166) issued by the Vétérinaire Cantonal (Vaud, Switzerland) and the Kanton Zürich Gesundheitsdirektion Veterinäramt (Zurich, Switzerland). Mice were housed up to 5 per cage in a temperature-controlled room with a 12hr light/dark cycle and ad libitum access to water and standard laboratory chow (Kliba, Nafag). For the experiments involving western diet, mice had ad libitum access to high-fat high-sugar food pellets (SAFE, Western HF Close TD88137 +0.15% Cholesterol).

The following transgenic mouse lines were used: Tg(Cnp-EGFP*)1Qrlu/J (JAX:026105), B6- Tmem117^tm^^1(tdTomato)Thor^/N^7^, B6.129-Olig2^tm^^1.1(cre)Wdr^/J (JAX:025567), B6.Cg-Tg(Plp1-cre/ERT)3Pop/J (JAX:005975).

### Cell lines GT1-7 cells

The mouse hypothalamic neuronal cell-line GT1-7^23^ was cultured in DMEM supplemented with 10% FBS, 5% horse serum and 1% penicillin-streptomycin on poly-L-Lysine coated plates. Cells were incubated at 37°C in a 5% CO2 atmosphere and passaged once per week up to 30 passages.

### MIN6B1 cells

The mouse pancreatic beta cell-line MIN6B1^24^ was cultured in DMEM supplemented with 15% fetal bovine serum (FBS), 71μM 2-mercaptoethanol and 1% penicillin-streptomycin. Cells were incubated at 37°C in a 5% CO2 atmosphere and passaged once per week up to 30 passages.

### HEK293T cells

The human kidney epithelial cell-line HEK293T^25^ was cultured in DMEM supplemented with 10% FBS and 1% penicillin-streptomycin. Cells were incubated at 37°C in a 5% CO2 atmosphere and passaged once per week up to 40 passages.

## METHOD DETAILS

### In vivo

#### Insulin-induced hypoglycemia, blood collection

Mice were food deprived for 6 hours (8am-2pm). At 12pm, they were placed in individual cages and glycemia was measured. At 1pm, glycemia was measured again and the mice were injected i.p. with either saline (baseline control) or insulin (0.8U/kg). At 2p.m. glycemia was measured again and 100- 120μl of blood were collected under isoflurane-induced general anesthesia by submandibular vein incision.

### Glucose tolerance test

Mice were food deprived for 12-14 hours before the test. Glucose was injected i.p in concentration of 2gr/kg. Glycemia was measured from tail blood taken at the indicated times after injection.

### Vagal nerve recordings

The firing rates of the thoracic branch of the vagal nerve along the carotid artery were recorded as previously described^26,27^. Unipolar parasympathetic activity was recorded in 5 h-fasted 8–10 weeks old mice. Recordings were performed for one and a half hour under isoflurane anesthesia [for 30 min during basal condition then for 1 h following i.p. insulin (0.8 U/kg) injection] using the LabChart 8 software (AD Instrument, Oxford, UK). Data were digitized with PowerLab 16/35 (AD Instrument, Oxford, UK). Signals were amplified 100 times and filtered using 100/1000 Hz band pass filter. Firing rate analysis was performed using LabChart 8.

### Estrus cycle monitoring

The estrus cycle phase was identified based on vaginal cytology samples after crystal violet staining^28^.

### Tamoxifen administration

Tamoxifen was diluted in corn oil at a concentration of 20mg/ml. Each mouse received an i.p injection of 100μl for 5 consecutive days.

### Ex vivo

#### Tissue collection

For immunofluorescence microscopy analysis, mice were transcardially perfused with 10ml ice-cold phosphate buffered saline (PBS: 137mM NaCl, 2.7mM KCl, 10mM Na2HPO4, 1.8mM KH2PO4) followed by 30ml ice-cold paraformaldehyde (PFA, 4%) in PBS. Then the brain was postfixed in 4% PFA O/N at 4◦C followed by 24-72 hours of cryopreservation in 30% sucrose in PBS at 4◦C. Tissues were frozen and stored at -80◦C.

For transmission electron microscopy analysis, mice were transcardially perfused with 10ml ice-cold 0.1M phosphate buffer (PB) followed by 40ml fresh cold fixative solution (2.5% glutaraldehyde in 0.1 M PB, pH 7.4). Then the brain was postfixed in the same fixative solution O/N at 4◦C followed by two washes with PB solution and slicing in 250μm-thick sagittal sections containing the *Corpus callosum* using a vibratome.

For RNA and protein isolation from the *Corpus callosum*, fresh brain tissue was sliced in 1mm coronal sections using a mouse brain slicing mold. Then the *Corpus callosum* was dissected using a scalpel from 2 consecutive sections (bregma ∼ 1 to -1), immediately quick frozen using dry ice and stored at -80◦C.

For a full phenotyping, mice were transcardially perfused with 10ml ice-cold PBS, followed by 30ml ice-cold paraformaldehyde (PFA, 4%) in PBS. The cadaver was then immersed in 10% buffered formalin for full fixation. All fixed animals were subjected to a full postmortem examination and gross assessment. Representative samples from all major organs were collected for histological examination.

### RNA extraction, cDNA synthesis and qRT-PCR

Total RNA was extracted from the *Corpus Callosum* samples using the QIAGEN RNeasy Mini kit based on manufacturer’s instructions. RNA was then reverse transcribed using M-MLV reverse transcriptase and the derived cDNA was subjected to a SYBR-Green based qRT-PCR for the quantification of the transcripts of interest. The primers used were: *Tmem117* recombined locus (FW:5’-CTTTCTTCATAAAAAGCCGGAAGGCATTAC-3’; RV:5’- GCCTGAAATATAAATATCGCAAGTGAGTGTGC-3’), *Xbp1* total (FW:5’- TGGCCGGGTCTGCTGAGTCCG -3’; RV:5’-ATCCATGGGGAGATGTTCTGG -3’), *sXbp1* (FW:5’-CTGAGTCCGAATCAGGTGCAG-3’; RV:5’-ATCCATGGGGAGATGTTCTGG-3’).

Expression data were normalized to the housekeeping gene *Gusb* (FW: 5’- CCACCAGGGACCATCCAAT-3’; RV: 5’-AGTCAAAATATGTGTTCTGGACAAAGTAA-3’).

### Protein extraction

Total protein from the same *Corpus Callosum* samples was isolated from the first column flow- through obtained during the RNA extraction. In brief, the flow-through was mixed with 9-volumes of -20◦C -cold methanol, incubated for 1 hour at -20◦C and then centrifuged at 4000rpm for 10 minutes.

The protein pellet was then diluted in RIPA buffer containing protease and phosphatase inhibitors.

### Proteomics

Samples were digested following the SP3 method^29^ using magnetic Sera-Mag Speedbeads (Cytiva 45152105050250, 50 mg/ml). Briefly, protein extracts (120 ug) precipitated with methanol were resuspended in SP3 buffer (2% SDS, 10mM DTT, 50 mM Tris, pH 7.5) and heated 10 min at 75°C. Proteins were then alkylated with 32mM (final) iodoacetamide for 45 min at RT in the dark. Beads were added at a ratio 10:1 (w:w) to samples, and proteins were precipitated on beads with ethanol (final concentration: 60 %). After 3 washes with 80% ethanol, beads were digested in 50ul of 100 mM ammonium bicarbonate with 2.4 ug of trypsin (Promega #V5073). After 1h of incubation at 37°C, the same amount of trypsin was added for an additional 1h of incubation. Supernatant were recovered, and acidified with formic acid (0.5% final concentration). To remove traces of SDS, two volumes of isopropanol, 1% TFA were added to the digests, and the samples were desalted on a strong cation exchange (SCX) plate (Oasis MCX; Waters Corp., Milford, MA) by centrifugation. After washing with isopropanol/1%TFA and 2% acetonitrile/0.1% FA, peptides were eluted in 200ul of 80% MeCN, 19% water, 1% (v/v) ammonia, and dried by centrifugal evaporation.

Aliquots (1/8) of samples were pooled and separated into 6 fractions by off-line basic reversed-phase (bRP) using the Pierce High pH Reversed-Phase Peptide Fractionation Kit (Thermo Fisher Scientific). The fractions were collected in 7.5, 10, 12.5, 15, 20 and 50% acetonitrile in 0.1 % triethylamine (∼pH 10). Dried bRP fractions were redissolved in 30 ul 2% acetonitrile with 0.5% TFA, and 3 ul were injected for LC-MS/MS analyses. Individual samples were resuspended in 1000ul of 2% acetonitrile,0.1% formic acid and 3 ul were injected.

LC-MS/MS analyses were carried out on a TIMS-TOF Pro (Bruker, Bremen, Germany) mass spectrometer interfaced through a nanospray ion source (“captive spray”) to an Ultimate 3000 RSLCnano HPLC system (Dionex). Peptides were separated on a reversed-phase custom packed 45 cm C18 column (75 μm ID, 100Å, Reprosil Pur 1.9 um particles, Dr. Maisch, Germany) at a flow rate of 250 nl/min with a 2-27% acetonitrile gradient in 93 min followed by a ramp to 45% in 15 min and to 90% in 5 min (total method time: 140 min, all solvents contained 0.1% formic acid). Identical LC gradients were used for DDA and DIA measurements.

For creation of the spectral library, data-dependent acquisitions (DDA) were carried out on the 6 bRP fractions sample pool using a standard TIMS PASEF method^30^ with ion accumulation for 100 ms for each survey MS1 scan and the TIMS-coupled MS2 scans. Duty cycle was kept at 100%. Up to 10 precursors were targeted per TIMS scan. Precursor isolation was done with a 2 Th or 3 Th windows below or above m/z 800, respectively. The minimum threshold intensity for precursor selection was 2500. If the inclusion list allowed it, precursors were targeted more than one time to reach a minimum target total intensity of 20’000. Collision energy was ramped linearly based uniquely on the 1/k0 values from 20 (at 1/k0=0.6) to 59 eV (at 1/k0=1.6). Total duration of a scan cycle including one survey and 10 MS2 TIMS scans was 1.16 s. Precursors could be targeted again in subsequent cycles if their signal increased by a factor 4.0 or more. After selection in one cycle, precursors were excluded from further selection for 60s. Mass resolution in all MS measurements was approximately 35’000.

The data-independent acquisition (DIA) used mostly the same instrument parameters as the DDA method and was as reported previously^31^. Per cycle, the mass range 400-1200 m/z was covered by a total of 32 windows, each 25 Th wide and a 1/k0 range of 0.3. Collision energy and resolution settings were the same as in the DDA method. Two windows were acquired per TIMS scan (100ms) so that the total cycle time was 1.7 s.

Raw Bruker MS data were processed directly with Spectronaut 16.3 (Biognosys, Schlieren, Switzerland). A library was constructed from the DDA bRP fraction data by searching the UNIPROT- SWISSPROT mouse proteome (www.uniprot.org) database of January 7th, 2022 and a contaminant database containing the most usual environmental contaminants and enzymes used for digestion^32^. For identification, peptides of 7-52 AA length were considered, cleaved with trypsin/P specificity and a maximum of 2 missed cleavages. Carbamidomethylation of cysteine (fixed), methionine oxidation and N-terminal protein acetylation (variable) were the modifications applied. Mass calibration was dynamic and based on a first database search. The Pulsar engine was used for peptide identification. Protein inference was performed with the IDPicker algorithm. Spectra, peptide and protein identifications were all filtered at 1% FDR against a decoy database.

Specific filtering for library construction removed fragments corresponding to less than 3 AA and fragments outside the 300-1800 m/z range. Also, only fragments with a minimum base peak intensity of 5% were kept. Precursors with less than 3 fragments were also eliminated and only the best 6 fragments were kept per precursor. No filtering was done on the basis of charge state and a maximum of 2 missed cleavages was allowed. Shared (non proteotypic) peptides were kept.

Peptide-centric analysis of DIA data was done with Spectronaut 15.7 using the library described above. Single hits proteins (defined as matched by one stripped sequence only) were kept in the Spectronaut analysis. Peptide quantitation was based on XIC area, for which a minimum of 1 and a maximum of 3 (the 3 best) precursors were considered for each peptide, from which the median value was selected. Quantities for protein groups were derived from inter-run peptide ratios based on MaxLFQ algorithm^33^. Global normalization of runs/samples was done based on the median of peptides.

Overall 97’900 peptide precursor ions were quantified in the dataset, mapped to 7’131 protein groups. 83’014 precursors (6’666 protein groups) had full profiles, i.e. were quantified in all samples. The average number of data points per LC peak was 6.2.

All subsequent analyses were done with an in house developed software tool (available on https://github.com/UNIL-PAF/taram-backend). Contaminant proteins were removed, and quantity values for protein groups generated by Spectronaut were log2-transformed. After assignment to groups, only proteins quantified in at least 4/4 samples in at least one group were kept. Missing values were imputed based on a normal distribution with a width of 0.3 standard deviations (SD), down- shifted by 1.8 SD relative to the median. Student’s T-tests were carried out among conditions, with Benjamini-Hochberg correction for multiple testing (adjusted p-value threshold <0.05). Imputed values were later removed. The difference of means obtained from the tests were used for 1D enrichment analysis on associated GO/KEGG annotations as described^34^. The enrichment analysis was also FDR-filtered (Benjamini-Hochberg, adjusted p-value <0.02).

### Quantification of circulating glucagon

Glucagon levels in the plasma were quantified by Glucagon ELISA (Mercodia, Cat# 10-1281-01), based on the manufacturer’s instructions.

### Immunofluorescent staining and microscopy

For mouse brain, 25 μm-thick tissue sections were washed with PBS for 5min at RT, incubated with blocking solution (2% normal goat serum + 0.3% Triton x-100 in PBS) for 1 hour at RT and then incubated O/N with primary antibody 1:250 (Rabbit anti-Tmem117, Novus, Cat# NBP1-94078) in blocking solution at 4◦C. Next, sections were washed with PBS three times for 10min, incubated with secondary antibody 1:500 (Donkey anti-Rabbit Alexa 647) in blocking solution for 2 hours at RT, washed again with PBS three times for 10min, incubated with DAPI (1:10000 in PBS) for 20min at RT, washed with PBS three times for 10min, allowed to dry and finally mounted using fluoromount (Sigma F4680). Images were captured with a Zeiss LSM 800 confocal microscope using the AiryScan mode.

### Transmission electron microscopy

Sections were stained (1% osmium tetroxide and 1.5% potassium ferrocyanide in 0.1M PB) for 1 hour followed by 1 hour incubation in 1% (wt/vol) osmium tetroxide in 0.1 M PB buffer. Samples were then stained for 1 hour in 1% Uranyl acetate in water before being dehydrated in a series of graded ethanol solutions (30%, 50%, 70%, 90%, 95%, 100% 3x). then sections were in 50/50 solution of ethanol/Durcupan resin for 1 hour followed by 100% Durcupan overnight. Next morning samples were put in fresh 100% resin and flat-embedded between aclar sheets. Samples were cured in oven at 60% for 24 hours. Trimmed blocks were used to cut ultrathin sections that were examined using a 120 kV TECNAI T12 electron microscope equipped with a Eagle camera (TFS).

### G-ratio analysis

The TEM images of the *Corpus Callosum* were imported in ImageJ and transformed into masks segmenting the myelin sheaths. A blinded experimenter outlined the inner and outer diameter for all axons that presented an intact circular morphology. G-ratio for each axon was defined as the ratio of the inner to the outer diameter.

### Histological examination

As part of the full histological phenotyping, formalin-fixed samples from all major organs were routinely trimmed and embedded in paraffin wax. Sections (2-4 µm) were prepared and stained with hematoxylin and eosin (HE) and histologically examined by a veterinary pathologist blinded to the grouping of the animals.

From the livers, one slice from the left lateral lobe was embedded into an OCT cryoblock using PrestoCHILL (Milestone Srl, Bergamo, Italy). A cryosection (8 µm) was prepared and routinely stained with Oil Red Orange (ORO) to visualize lipid accumulation. From the remaining liver, 2-3 slices were trimmed and routinely embedded in paraffin. Consecutive sections (2-4 μm) were prepared and stained with haematoxylin and eosin (HE) for general histological evaluation and subjected to the periodic acid-Schiff (PAS) reaction to assess glycogen accumulation in hepatocytes. The liver sections were histologically assessed by a veterinary pathologist blinded to the grouping of the animals, to assess nonalcoholic steatohepatitis (NASH), i.e. hepatocellular fat accumulation, inflammation, liver cell injury, and grading the changes according to the total Kleiner score^35^ which was modified for the assessment of mouse liver: (a) the presence of microvesicular steatosis (which is consistently seen in the normal mouse liver) was noted but not further acknowledged in the scoring; (b) scattered random small leukocyte aggregates consistent with the mild extramedullary hematopoesis frequently observed incidentally also in normal mouse livers were not considered as lobular inflammation.

### In vitro

#### Intracellular calcium quantification with Fluo-4

MIN6B1 cells were plated in 96-well black tissue culture plate (Costar, REF# 3603) at a density of 21’000 cells/well. After 48 hours cells were transfected with the pRRL-PGK-Tmem117-flag-puroR plasmid or the pRRL-PGK-GFP-puroR plasmid (100ng/well) with lipofectamine 3000 (Thermo Fisher Scientific, Cat# L3000001). 48 hours after transfection, cells were incubated in KRBH buffer (120mM NaCl, 4mM KH2PO4, 20mM Hepes, 1mM MgCl2, 1mM CaCl2, 5mM NaHCO3) supplemented with 0.5% BSA and 2.5mM glucose [low glucose buffer (LG)] for 2 hours. Then LG buffer was replaced by LG+Fluo-4 (2μM, Thermo Fisher Scientific, Cat# F14201)+probenecid (2mM, Selleck Chemicals, Cat# S4022) for 1 hour. Finally, LG+Fluo-4+probenecid was replaced by LG or LG+KCl (40mM) and 5 minutes later fluorescent signal (480nm excitation/520nm emission) was quantified using a TECAN plate reader (SAFIRE II).

For GT1-7 cells, 40’000 cells were plated per well in Poly-L-Lysine coated 96-well black tissue culture plate (Costar, REF# 3603). After that the protocol was exactly as described above for MIN6B1 cells.

### Intracellular calcium quantification with Fura red

HEK293T cells were plated in 12-well tissue culture plate (Falcon, REF# 353225) containing glass coverslips at a density of 300’000 cells/well. After 24 hours cells were co-transfected with the pcDNA3.1/TMEM117-eGFP (1.3μg/well) and the NCX1_pcDNA3.1/Zeo (0.7 μg/well) plasmids with PEI (7.5mM). Twenty-four hours after transfection the medium was supplemented with zeocin (100 μg/ml) for the selection of the NCX1 expressing cells.Forty-eight hours after transfection, cells were incubated in complete medium+zeocin+PowerLoad (1X, Thermo Fisher Scientific, Cat# P10020)+Fura red (5μM, Thermo Fisher Scientific, Cat# F3021)±SEA0400 (1μM, MCE, Cat# HY- 15515) for 1 hour. Then the medium was removed, cells were washed twice with PBS, fixed with 4% PFA, washed again with PBS and the coverslips were mounted on microscopy slides using fluoromount (Sigma F4680). Images were captured using a Zeiss LSM800 confocal microscope (excitation/emission wavelengths were 471/670 and 435/660 for unbound and Ca^2+^-bound forms, respectively. The detection window was set at 640-700 in both cases). Cell segmentation and intensity analysis was performed using QuPath by a blinded experimenter.

### Insulin secretion assay

MIN6B1 cells were plated in 12-well tissue culture plate (Falcon, REF# 353225) at a density of 400’000cells/well. After 48 hours cells were transfected with the pRRL-PGK-Tmem117-flag-puroR plasmid or the pRRL-PGK-GFP-puroR plasmid (2μg/well) with lipofectamine 3000 (Thermo Fisher Scientific, Cat# L3000001). Forty-eight hours after transfection, cells were incubated in KRBH buffer (120mM NaCl, 4mM KH2PO4, 20mM Hepes, 1mM MgCl2, 1mM CaCl2, 5mM NaHCO3) supplemented with 0.5% BSA and 2.5mM glucose [low glucose buffer (LG)] for 2 hours. Then LG buffer was replaced by LG+KCl (40mM) and 5 minutes later the supernatant was collected for quantification of the secreted insulin by ELISA (Mercodia, Cat# 10-1247-10). The remaining supernatant was removed, and cells were lysed in TETG buffer (20mM Tris-HCl, 1% Triton X-100, 10% glycerol, 137mM NaCl, 2mM EGTA) for quantification of insulin content and further protein analysis.

### Western blot

For the quantification of the Tmem117 protein levels in MIN6B1 cells used for the insulin secretion assay, the protein content of the samples was quantified by BCA assay. 20μg of total protein per sample were loaded on a 4-20% MP TGX Stain-Free premade Gel (bio-rad, Cat# 4568095). Proteins were transferred onto a nitrocellulose membrane, incubated with blocking solution [3% milk in PBS- T (PBS + 0.1% Tween-20)] for 1 hour at RT, washed with PBS-T 3 times for 10 minutes and incubated O/N at 4◦C with primary antibodies (Rabbit anti-Tmem117, Novus, Cat# NBP1-94078 and Rabbit anti-actin, Sigma, Cat# A2066) at a dilution of 1:100 and 1:1000, respectively, in PBS-T. Then membranes were washed with PBS-T 3 times for 10 minutes, incubated for 1 hour at RT with horseradish peroxidase (HRP) conjugated secondary antibody (Anti-Rabbit HRP, Amersham, Cat# NA9340) at a dilution of 1:10000 in blocking solution, washed again with PBS-T 3 times for 10 minutes, incubated for 1 minute with enhanced chemiluminescence (ECL, Amersham) buffer and the HRP mediated chemiluminescence signal was detected/imaged using the Fusion FX6 Spectra imaging platform (VILBER). The derived images were analyzed for band intensity using imageJ. The Tmem117 signal of each sample was normalized to the β-actin signal. Data are presented as percent of the control non-transfected cells.

For the quantification of the NCX1 protein levels, HEK293T cells were plated in 12-well tissue culture plate (Falcon, REF# 353225) at a density of 300’000 cells/well. After 24 hours cells were co-transfected with the pcDNA3.1/TMEM117-eGFP or pcDNA3.1/eGFP control (1.3μg/well) and the NCX1_pcDNA3.1/Zeo (0.7 μg/well) plasmids with PEI (7.5mM). Twenty-four hours after transfection the medium was supplemented with zeocin (100 μg/ml) for the selection of the NCX1 expressing cells. Forty-eight hours after transfection medium was removed, cells were washed twice with PBS and then lysed in RIPA buffer containing protease and phosphatase inhibitors. The protein content of the samples was quantified by BCA assay. 10μg of total protein per sample were loaded on a 4-20% MP TGX Stain-Free premade Gel (bio-rad, Cat# 4568095). Proteins were transferred onto a nitrocellulose membrane, incubated with blocking solution [3% milk in PBS-T (PBS + 0.1% Tween- 20)] for 1 hour at RT, washed with PBS-T 3 times for 10 minutes and incubated O/N at 4◦C with primary antibody (Mouse anti-NCX1, Thermo Fisher Scientific, Cat# MA3-926) at a dilution of 1:1000 in PBS-T. Then membranes were washed with PBS-T 3 times for 10 minutes, incubated for 1 hour at RT with horseradish peroxidase (HRP) conjugated secondary antibody (Anti-Mouse HRP, Amersham, Cat# NA931) at a dilution of 1:10000 in blocking solution, washed again with PBS-T 3 times for 10 minutes, incubated for 1 minute with enhanced chemiluminescence (ECL, Amersham) buffer and the HRP mediated chemiluminescence signal was detected/imaged using a LAS-1000 imaging system from Fuji. The derived images were analyzed for band intensity using imageJ. The NCX1 signal of each sample was normalized to the total protein (Pierce Reversible Protein Stain Kit). Data are presented as percent of the control.

### Immunofluorescent staining and microscopy

HEK293T cells were plated in 12-well tissue culture plate (Falcon, REF# 353225) containing glass coverslips at a density of 300’000 cells/well. After 24 hours cells were co-transfected with the pcDNA3.1/TMEM117-eGFP (1.3μg/well) and the NCX1_pcDNA3.1/Zeo (0.7 μg/well) plasmids with PEI (7.5mM). Twenty-four hours after transfection the medium was supplemented with zeocin (100 μg/ml) for the selection of the NCX1 expressing cells. Forty-eight hours after transfection the medium was removed, cells were washed twice with PBS, fixed with 4% PFA, washed again with PBS, incubated with blocking solution (2% normal goat serum + 0.3% Triton x-100 in PBS) for 1 hour at RT and then incubated O/N with primary antibody 1:500 (Mouse anti-NCX1, Thermo Fisher Scientific, Cat# MA3-926) in blocking solution at 4◦C. Next, coverslips were washed with PBS three times for 10 minutes, incubated with secondary antibody 1:500 (Donkey anti-Mouse Alexa 647) in blocking solution for 2 hours at RT, washed again with PBS three times for 10 minutes, incubated with DAPI (1:10000 in PBS) for 20 minutes at RT, washed with PBS three times for 10 minutes, allowed to dry and finally mounted using fluoromount (Sigma F4680). Images were captured using a Zeiss LSM800 confocal microscope. Cell segmentation and intensity analysis was performed using QuPath by a blinded experimenter.

### Proximity ligation assay

HEK293T cells were plated in 12-well tissue culture plate (Falcon, REF# 353225) containing PDL-coated glass coverslips at a density of 200’000 cells/well. After 24 hours cells were co-transfected with the pcDNA3.1/TMEM117-eGFP or pcDNA3.1/eGFP control (1.3μg/well) and the NCX1_pcDNA3.1/Zeo (0.7 μg/well) plasmids with PEI (7.5mM). 24 hours after transfection the medium was supplemented with zeocin (100 μg/ml) for the selection of the NCX1 expressing cells. 48 hours after transfection the medium was removed, cells were washed twice with PBS, fixed with 4% PFA and washed again twice with PBS. Then the PLA was performed using the Duolink II kit (Merck) according to manufacturer’s instructions. In brief, the cells were permeabilized with 0.2% triton in PBS for 15 minutes at RT and incubated O/N with primary antibodies 1:500 (Rabbit anti-Tmem117, Novus, Cat# NBP1-94078; Mouse anti-NCX1, Thermo Fisher Scientific, Cat# MA3-926) in 10% NGS at 4◦C. Then the coverslips were washed four times for 5 minutes in PBS and incubated for 1 hour at 37◦C with the PLA probes (in a pre-heated humidity chamber). Then coverslips were washed twice for 5 minutes with pre-warmed Duolink II Wash Buffer A and incubated for 30 minutes at 37◦C with the ligation solution. Then coverslips were washed twice for 5 minutes with pre-warmed Duolink II Wash Buffer A and incubated for 1 hour and 40 minutes at 37◦C with the amplification solution. Then the coverslips were washed twice for 10 minutes with pre-warmed Duolink II Wash Buffer B and a 3^rd^ time for 1 minute with pre-warmed Duolink II Wash Buffer B diluted 1:100 in ddH2O. Finally, the coverslips were incubated with DAPI (1:10000 in PBS) for 10 minutes, washed twice for 2 minutes with PBS and mounted on microscopy slides using fluoromount (Sigma F4680). Images were captured using a Zeiss LSM800 confocal microscope. The number of puncta/particles per image were quantified using imageJ.

### Constructs / Cloning

The pcDNA3.1/TMEM117-eGFP plasmid containing the *H.sapiens Tmem117* CDS tagged with an eGFP in the C-terminus was provided by the Van der Goot group^36^. The pRRL-PGK-Tmem117-flag-puroR plasmid containing the *M.musculus Tmem117* CDS was constructed in the Thorens group by Dr. Simon Quenneville. The NCX1_pcDNA3.1/Zeo plasmid containing the *H.sapiens Slc8a1* CDS was generated by GenScript.

## BIOINFORMATIC ANALYSIS

### Protein network analysis/ STRING

For the analysis of the differentially abundant proteins detected in the proteomic analysis of the *Corpus callosum* we imported the list of the proteins that had a p value < 0.05 into STRING to observe the protein-protein interaction networks and perform functional enrichment analysis. To obtain the functional enrichment graph included in Fig 4E, KEGG Pathways was selected as the plotting category with the following specifications: group terms by similarity ≥ 0.8; sort terms by signal; no merge of the rows by term similarity; maximum FDR ≤ 0.05; minimum signal ≥ 0.01; minimum strength ≥ 0.01; minimum count in network 2. To obtain the protein-protein networks presented in Fig 4F, a k-mean clustering analysis was performed using the same online tool with k=40 and the nodes composed of ≥ 4 proteins were included alongside the description/assigned term.

### Differential expression analysis

For the differential expression analysis between *Tmem117*-enriched and -non enriched cells, we analyzed 3 datasets: cells of the human brain white matter^12^, mouse cells from 20 different organs^10^ and cells of the sea anemone^13^. For the human dataset, cells with mitochondrial gene content exceeding 2.59% were excluded. The remaining data were normalized using Seurat’s “LogNormalize” method and were then divided into two groups: cells showing high levels of *Tmem117* expression (>3 LogNormalized counts) and cells with low *Tmem117* expression (<1 LogNormalized counts). Differential gene expression analysis between these two groups was performed using the Seurat package. A similar workflow was applied to the Tabula Muris dataset, with the only modification being a higher mitochondrial gene content cut-off of 5%. Lastly, for the *N. vectensis* dataset the cells were again separated to *v1g11324*-high (>1 molecule/1000 UMIs) and *v1g11324*-low (<1 molecule/1000 UMIs), and then differential gene expression analysis between these two groups was performed using the Seurat package. The original code used is available from the lead contact upon request.

## QUANTIFICATION AND STATISTICAL ANALYSIS

All graphs and statistical analysis were generated using Prism software (GraphPad Prism 10), further details regarding sample size and the statistical analysis used in each case can be found in the corresponding figure legends. In brief, for experiments concerning comparisons between two groups on a single independent variable (Fig 4B-C, 5J, 5L) we used unpaired two-tailed t test. For experiments concerning comparisons among three groups on a single independent variable (Fig 5C and Suppl Fig 5A-C) we used one-way ANOVA with post hoc tests. For two-factor designs concerning repeated measures (Fig 2B, D-F, Suppl Fig 2B-D, Fig 3C-D, H-I and Suppl Fig 3B-C) we used two-way ANOVA RM with post hoc tests. For two-factor designs concerning non-repeated measures (Fig 3M, Suppl Fig 3D-E, I and Fig 5D) we used two-way ANOVA with post hoc tests. For three-factor designs concerning repeated measures (Fig 3K-L and Suppl Fig 3G-H) we used three-way ANOVA RM. For experiments concerning multiple comparisons between two groups on a single independent variable (Fig 2G) we used multiple unpaired two-tailed t test. Lastly, for experiments concerning correlation of two independent variables (Fig 5E-F, H and Suppl Fig 5E) we used Pearson’s r correlation analysis.

## Notes

### Competing Interest Statement

The authors have declared no competing interest.

